# Practical Speedup of Bayesian Inference of Species Phylogenies by Restricting the Space of Gene Trees

**DOI:** 10.1101/770784

**Authors:** Yaxuan Wang, Huw A. Ogilvie, Luay Nakhleh

## Abstract

Species tree inference from multi-locus data has emerged as a powerful paradigm in the post-genomic era, both in terms of the accuracy of the species tree it produces as well as in terms of elucidating the processes that shaped the evolutionary history. Bayesian methods for species tree inference are desirable in this area as they have been shown to yield accurate estimates, but also to naturally provide measures of confidence in those estimates. However, the heavy computational requirements of Bayesian inference have limited the applicability of such methods to very small data sets.

In this paper, we show that the computational efficiency of Bayesian inference under the multispecies coalescent can be improved in practice by restricting the space of the gene trees explored during the random walk, without sacrificing accuracy as measured by various metrics. The idea is to first infer constraints on the trees of the individual loci in the form of unresolved gene trees, and then to restrict the sampler to consider only resolutions of the constrained trees. We demonstrate the improvements gained by such an approach on both simulated and biological data.

## 1 Introduction

Species tree inference under the multispecies coalescent (MSC) model accounts for gene tree heterogenity that arises due to incomplete lineage sorting (Tajima, 1983). This model has gained much attention in the years since the first inferential methods to implement it were developed (Takahata *et al*., 1995; Yang, 2002; Degnan and Rosenberg, 2009). A wide array of methods that assume or are inspired by the MSC have been devised (Liu *et al*., 2010; Liu and Yu, 2011; Mirarab *et al*., 2014; Chifman and Kubatko, 2014; Wang and Nakhleh, 2018), including the Bayesian methods of Ogilvie *et al*. (2017); Flouri *et al*. (2018); Zhu *et al*. (2018). The MSC was recently extended to the multispecies network coalescent to account for reticulation (in addition to incomplete lineage sorting; see Yu *et al*., 2014) and Bayesian methods for inference under this model have been devised (Wen *et al*., 2016; Wen and Nakhleh, 2017; Wen *et al*., 2018; Zhu *et al*., 2018; Zhang *et al*., 2017).

The power of Bayesian methods lies in their ability to incorporate prior knowledge, infer values of parameters beyond the tree topology, and provide measures of confidence in the inference based on the posterior that they sample (Huelsenbeck *et al*., 2001). However, for these Bayesian methods to approximate the true posterior distribution, they demand significant computational resources, an issue that has thus far limited their applicability in terms of both the number of taxa and number of loci in the data set (Ogilvie *et al*., 2016). This is why incomplete lineage sorting aware methods that have been proven to be statistically consistent under the MSC and, at the same time very efficient computationally, are used to infer large-scale species trees (Mirarab *et al*., 2014; Liu *et al*., 2010; Liu and Yu, 2011; Chifman and Kubatko, 2014). However, these methods focus almost exclusively on the species tree topology and provide neither accurate information on other parameters, such as divergence times and population sizes, nor confidence intervals for their inferences. The question we address in this paper is: Can the convergence of Bayesian methods be improved in practice without sacrificing the accuracy of the information they provide?

A rich body of literature exists on the development of methods for statistical inference outside phylogenetics, much of which has been adopted by Bayesian phylogenetic methods. The ubiquitous Markov chain Monte Carlo (MCMC) arose from nuclear weapons research (Robert and Casella, 2011), and is the basis for tree and network inference in MrBayes, *BEAST, PhyloNet and other software tools. The efficiency of MCMC for phylogenetics has been improved with the development of new MCMC proposals (e.g. Höhna and Drummond, 2011; Zhang *et al*., 2019; Yang and Rodríguez, 2013), including proposals designed to improve the mixing of MSC models (e.g. Yang and Rannala, 2014; Rannala and Yang, 2017; Jones, 2017).

Metropolis coupling to accelerate MCMC was developed for the inference of spatial statistics (Geyer, 1991), and has been implemented in various phylogenetics software (Ronquist *et al*., 2012; Bouckaert *et al*., 2019; Wen *et al*., 2018). Variational Bayes is a radically different approach which fits parametric distributions to model parameters, unlike MCMC which is non-parametric. Variational Bayes was originally developed for graphical models (Attias, 1999), and has recently been applied to compute posterior distributions and marginal likelihoods of phylogenetic trees (Zhang and Matsen, 2019; Fourment and Darling, 2019).

All of these methods were developed decades before their adoption for phylogenetic inference because tree and network space is far more complex than the typical multidimensional parameter space. The number of unrooted or rooted trees grows superexponentially with the number of taxa (Felsenstein, 1978), and is for all practical purposes infinite when the number of taxa is large. Multilocus MSC inference embeds gene trees within a species tree, with the constraint that between-species coalescent events must take place earlier in time than the most recent common ancestor (MRCA) time of the involved species. This multiplies the complexity of the inference problem by increasing the number of trees to infer, and because the probability distributions of node heights for different trees are not independent.

Rather than trying to adapt an algorithm developed for other fields of natural sciences or mathematics, we have developed a heuristic method that specifically applies to the problem of multilocus MSC inference. The heuristic method constrains the space of gene tree topologies to allow for faster convergence and, consequently, analyses of larger data sets. The idea behind our approach is very simple: A set of constraints in the form of a tree which is usually less than fully resolved is estimated independently for each individual locus, and then MCMC walks in the portion of the tree space that is consistent with these constraints.

In other words, the MCMC sampler considers only gene trees that are consistent with the constraints on the individual loci. Using simulated data under a variety of conditions and employing several metrics for assessing performance, we demonstrate that this simple approach results in computational improvements relative to unconstrained Bayesian MCMC without sacrificing accuracy. We then analyze a biological data set and show that the new approach enables analyses that had before necessitated dividing the data set into smaller ones.

Our work presents an approach for improving the computational requirements of Bayesian inference of species phylogenies. The constraints on the individual loci can be obtained in various ways and the proposals that satisfy these constraints can be derived in multiple ways as well. In this work, we implemented one specific method for obtaining the constraints and a standard set of proposals that satisfy them. As this approach can be adopted by any Bayesian species phylogeny inference method, both of these components can be further modified to achieve even further improvement to the computational requirements of Bayesian inference.

## 2 New Approach

A Bayesian formulation of the multi-locus species tree inference problem is to estimate the posterior distribution over species tree topologies, population sizes and species divergence times from multiple sequence alignments given a model which includes at least some demographic function (e.g. constant population sizes) and substitution process (e.g. Jukes and Cantor, 1969).

Here we use *S* to represent the species tree, Θ to represent the population sizes and divergence times, and *X* to represent the multiple sequence alignments. Inspired by Rannala and Yang (2017), we can formulate the model as

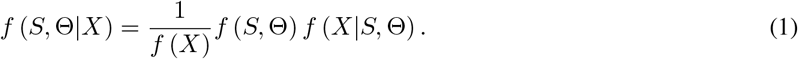

In this formulation, *f* (*S*, Θ|*X*) is the posterior probability of the species tree topology and associated parameters, *f* (*X*) is the marginal likelihood, *f* (*S*, Θ) is the prior on the species tree topology and associated parameters, and *f* (*X*|*S*, Θ) is the likelihood. To calculate the likelihood we must integrate over gene trees *G* and substitution model parameters *ψ*:

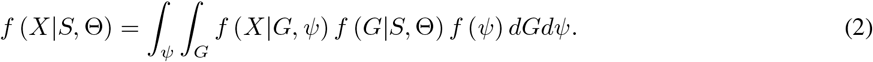

In the above formulation, *f* (*X*|*G*, *ψ*) is the phylogenetic likelihood and substitution model parameters, *f* (*G*|*S*, Θ) is the coalescent likelihood, and *f* (*ψ*) is the prior for the same parameters. Note that the coalescent and phylogenetic likelihoods are functions of a gene tree, and are calculated over the space of all possible gene trees. When using MCMC, we only need a density proportional to the posterior probability, so the marginal likelihood can be omitted.

Furthermore, under the common assumption of recombination-free, unlinked loci, the likelihood can be derived from the product of integrations for each locus:

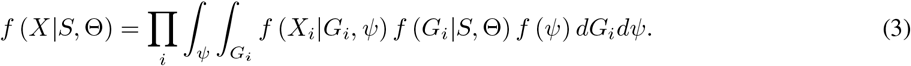

where *X_i_* is the multiple sequence alignment of the *i*th locus in the data set and *G_i_* is the gene tree sampled at locus *i*.

As the integration in Equation (3) cannot be derived analytically, MCMC sampling algorithms are often employed to obtain samples from the posterior distribution and approximate it based on those samples. Due to the scaling problems of MCMC inference (Ogilvie *et al*., 2016), current algorithms to approximate Equation (3) become computationally infeasible for larger data sets, getting stuck in the peaks and troughs of the posterior distribution and requiring extremely large numbers of iterations to converge.

Our approach to tackle the computational challenge works as follows. For each sequence alignment *X_i_*, maximum likelihood with bootstrapping is run to obtain a set of gene trees from which a majority-rule consensus tree with a pre-specified support threshold *x* is built. For example, for *x* = 90, a majority-rule consensus tree is built where only clades that appear in at least 90% of the bootstrap trees are included. We denote this majority-rule consensus tree by *C_i_* (we use the value of *x* explicitly in the naming only when it is not clear from the context) and call it a constraint gene tree, or CGT. our approach now samples according to Equation (3) with one difference: The integration is taken over gene trees that are consistent with, i.e., refinements of, their respective *C_i_* constraints. For example, if *C_i_* = (((*A, B*),*C*), (*D, E, F*)), the sampler considers gene tree *G_i_* = (((*A, B*), *C*), ((*D, E), F*)) as it is a refinement of *C_i_*, but does not consider gene tree *G_i_* = (((*A,B*), (*C,D*)), (*E, F*)) as it is not a refinement of *C_i_*. This concept is illustrated in Figure 1.

**Figure 1:**
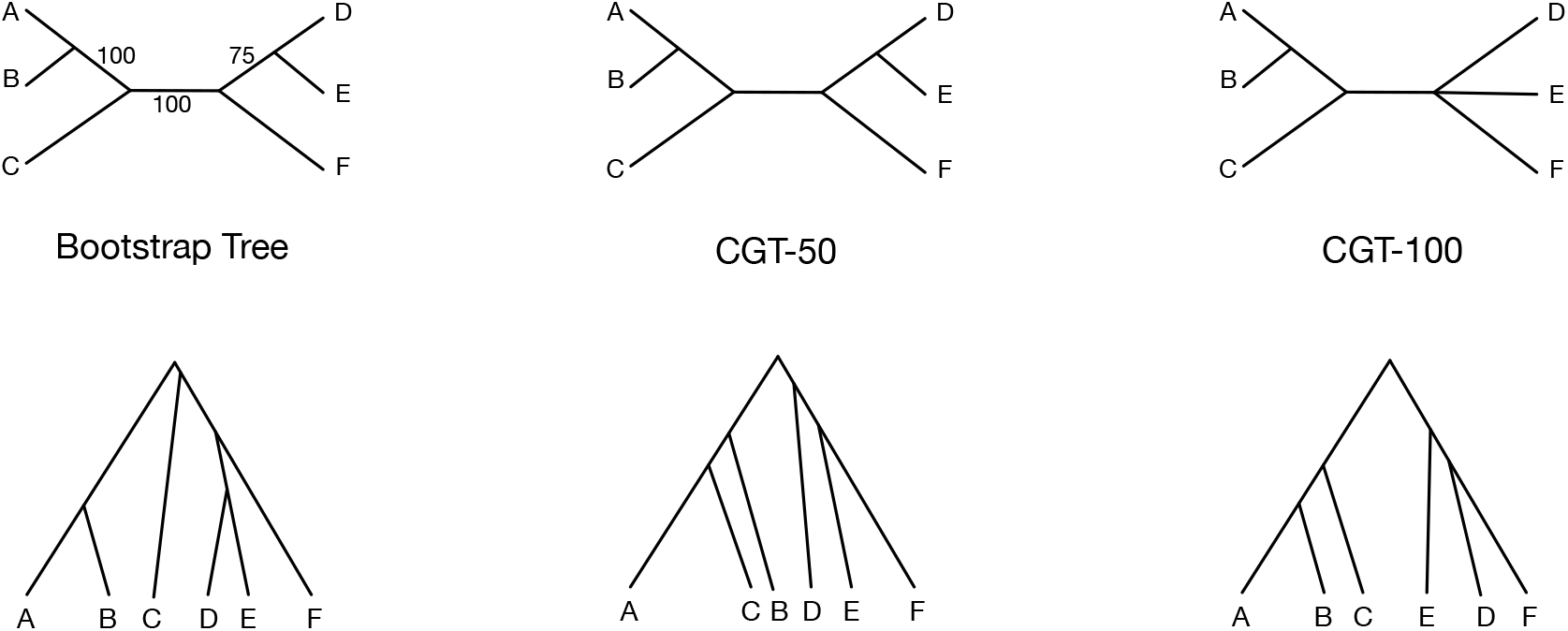
Constraint trees, their resolutions, and acceptable moves. Bootstrap tree and CGTs under the consensus threshold of 50 (CGT-50) and 100 (CGT-100) are shown in the first row. In the second row, three possible proposed gene trees are provided. The left tree is acceptable given the constraints CGT-50 and CGT-100. The middle tree is not acceptable given the constraints CGT-50 or CGT-100. The right tree is acceptable given the constraint CGT-100 but not CGT-50.

We implemented this restricted sampling of gene tree topologies by comparing proposed topologies with the constraint trees, and rejecting incompatible proposals. Future implementations may be made even more efficient by only proposing compatible topologies.

Observe that if *C_i_* is a star phylogeny (a tree that has no internal branches), then the method is effectively sampling according to Equation (3), whereas if *C_i_* is fully resolved (a binary tree), then the sampler fixes the gene tree topology for locus *i* and only samples its parameters. It is important to note that *C_i_* imposes only topological constraints; that is, *C_i_* has no branch lengths. Furthermore, we take *C_i_* to be unrooted, so that the sampler is allowed to sample the roots of the gene trees.

The posterior of species trees includes a “long tail” region of model trees where the likelihood of each tree is very small but the number of trees within such region is very large. Unfortunately, as the scale of the data set increases, the autocorrelation time of the MCMC chain increases dramatically (Ogilvie *et al*., 2016). The motivation of our approach is that by constraining the gene trees, the sampler can avoid the long tail and have better mixing. While utilizing these constraints necessarily means that the sampler is not sampling from the same posterior distribution as an unconstrained version of the sampler (except for the case where the constraints are star phylogenies), we demonstrate below that this has very little impact on the accuracy of the sampler in practice.

Hereafter we write CMCMC to denote constrained MCMC according to our new approach and UMCMC to denote the unconstrained version of MCMC. We also write CMCMC-*x*, where *x* is a value between 50 and 100, to denote the use of CMCMC with support threshold *x* in the majority-rule consensus tree, or a value of 0 to denote use with the maximum likelihood tree.

Parameter values must be initialized somehow at the beginning of each MCMC chain. For both CMCMC and UMCMC we initialized gene trees by inferring the maximum likelihood (ML) tree for each locus in RAxML (Stamatakis, 2014). We used the topology of the ML tree, and for each internal node used the maximum distance from that node to any descendant leaf as the node height. Although there is a chance that the ML tree is incompatible with the constraint tree due to the stochastic nature of bootstrapping, this is probably very uncommon as we did not encounter this in any of our analyses.

## 3 Results and Discussion

A simulation study was carried out to comprehensively analyze the performance of CMCMC and UMCMC. We varied the simulation data set along three dimensions: evolutionary scenarios of species, the number of loci and the number of taxa. When we focused on one dimension, the other two dimensions were fixed. More details are provided in the Materials and Methods section. To simulate different scenarios of complexity and signal in the data, we varied evolutionary time scales and population sizes in four categories:

- “OH”: old divergence times and high population size;
- “OL”: old divergence times and low population size;
- “YH”: young divergence times and high population size; and,
- “YL”: young divergence times and low population size.

To further examine the performance of each method, we varied the number of loci (10, 20, 40) for the YH condition while fixing the number of taxa as 16. We also varied the numbers of taxa (16, 32, 48) for the YH condition and fixed the number of loci as 10. Unless otherwise stated, there are 10 replicates for each condition.

### 3.1 CGTs improve the convergence of MCMC

The ability to converge within a reasonable time is a key metric to evaluate the performance of an MCMC sampler. An effective sample size (ESS) of at least 200 is used as a threshold for convergence in the popular MCMC diagnostic and analysis tool Tracer (Rambaut *et al*., 2018). In this work, we target the same convergence standard for continuous parameters including the posterior probability, likelihood, prior probability, coalescent likelihood, tree height and population size. We terminated any chain still running after 72 hours.

Figure 2(a) shows that decreasing the consensus threshold enables convergence for low population size conditions, which is impossible for UMCMC within 72 hours. We also show the improved convergence of CMCMC as the number of loci increases in Figure S1 and as the number of taxa increases in Figure S2.

**Figure 2:**
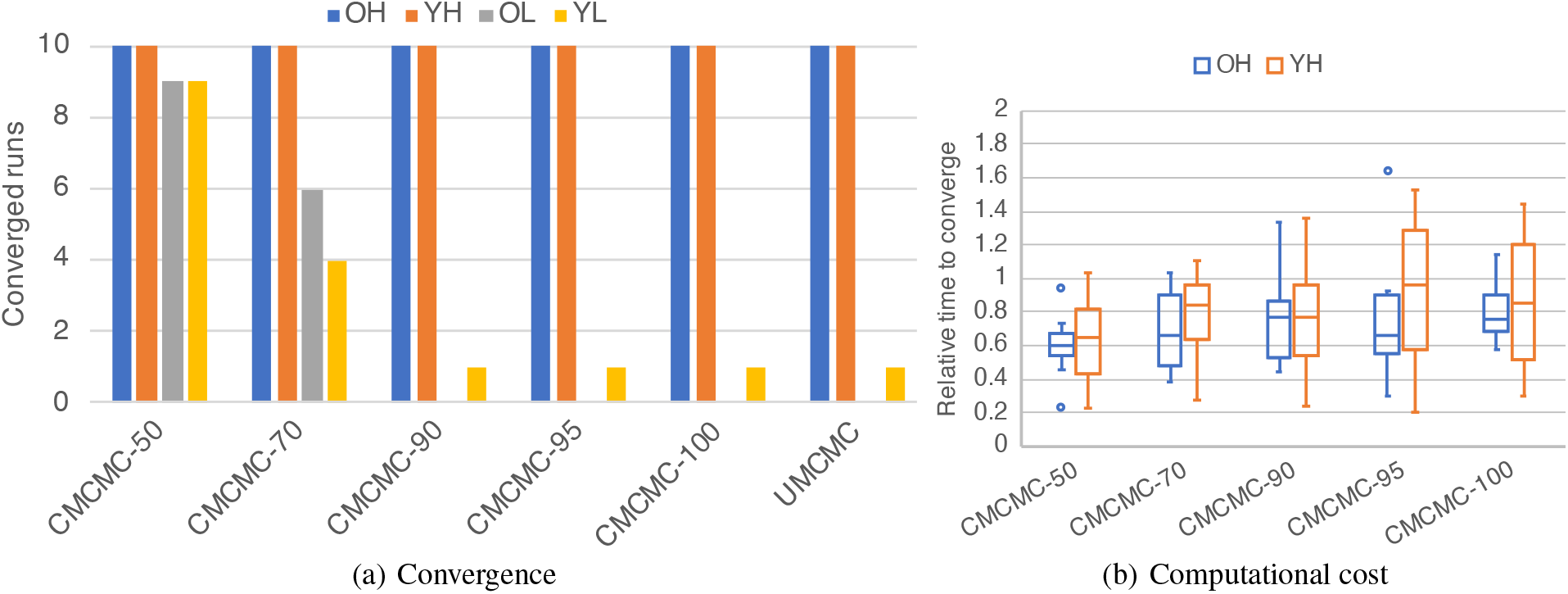
Convergence and efficiency of CMCMC and UMCMC. (a) Convergence of samplers with or without constraint gene trees. Different samplers are shown on the x axis and the y axis shows the number of data sets (out of 10) on which the sampler converged, (b) Ratios of iterations required for convergence using CMCMC compared with UMCMC. Values above 1 are faster using UMCMC, below 1 are faster using CMCMC.

When MCMC chains are able to converge, CMCMC reduces the number of iterations required for convergence into less than half that of UMCMC under various evolutionary parameters as shown in Figure 2(b). Furthermore, CMCMC took fewer iterations than UMCMC to converge for different numbers of loci and different numbers of taxa (Figure S3 and Figure S4).

### 3.2 CMCMC and UMCMC derive similar posterior distributions

The ultimate goal of phylogenetic inference problem with Bayesian sampling is to approximate the posterior distribution of the true species tree. One way to verify the posterior distribution is to compare the average standard deviation of split frequencies (ASDSF; Lakner *et al*., 2008) of the 95% credible set. Note that the 95% credible set or interval of a well calibrated Bayesian method covers the true value in 95% of cases. The smaller the ASDSF is, the more similar the species tree distributions are. A threshold of 0.01 on the ASDSF is commonly used to assess the convergence of two chains. An ASDSF value below 0.01 is taken to indicate that the chains are likely to be sampling from the same underlying distribution (for examples, see Stunžėnas *et al*., 2011; Mazza *et al*., 2016; Stensvold *et al*., 2011).

Figure 3 shows the ASDSF between the CMCMC and UMCMC chains for different evolutionary scenarios, different numbers of loci, and different numbers of taxa. To compare the ASDSF, the MCMC chains must be converged. However, although CMCMC has better convergence ability than UMCMC, we only compare the ASDSF when all MCMC chains can converge. For all simulated scenarios in Figure 3(a), the ASDSF values of CMCMC and UMCMC in most replicates are below 0.01. As we increased the number of loci and taxa, all ASDSF interquartile ranges (IQRs) fell below 0.01, except for CMCMC-50 for which the top of the range could be slightly above 0.01, as shown in Figure 3(b) and Figure 3(c).

**Figure 3:**
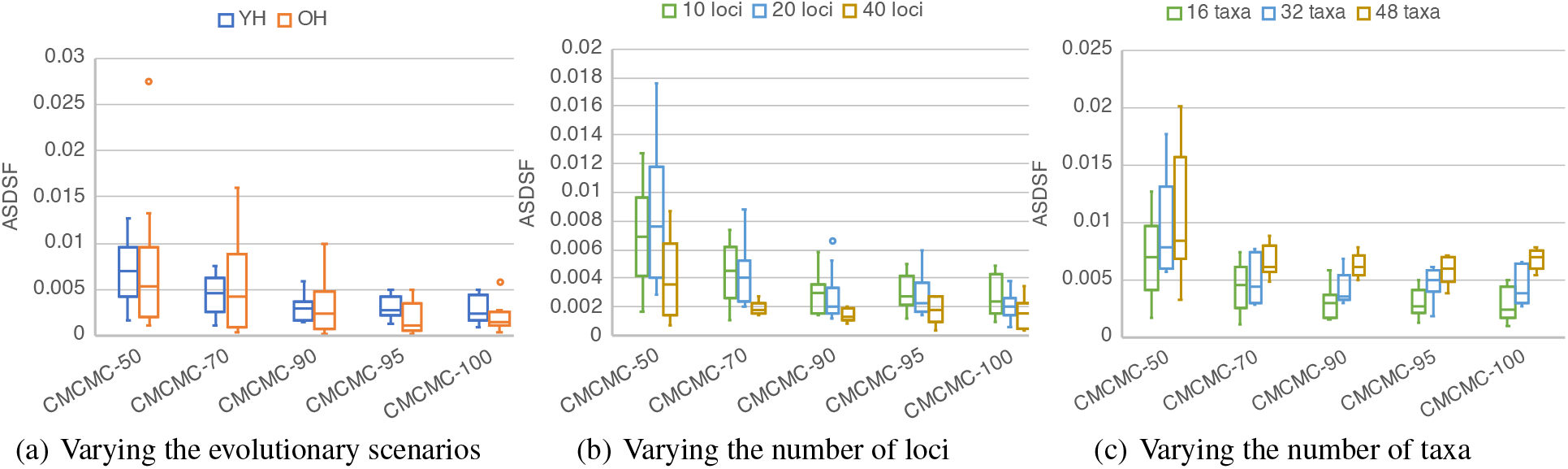
ASDSF between the CMCMC and UMCMC chains. The X axis lists CMCMC samplers with different support thresholds and the y axis shows the ASDSF between each CMCMC method and UMCMC. (a) ASDSF values when varying the divergence times fixing the number of taxa and loci as 16 and 10. Only ‘YH’ and ‘OH’ are shown because UMCMC cannot converge in ‘YL’ and ‘OL’ scenarios. (b) ASDSF values when varying the number of loci, restricted to the ‘YH’ scenario. There are 10 replicates for 10- and 20-locus data sets. For 40-locus data sets, results are shown for the 6 out of 10 replicates where all methods converged within 20 days. (c) ASDSF values when varying the number of taxa, restricted to the ‘YH’ scenario. There are 10 replicates for 16-, 32- and 48-taxon data sets.

As the number of loci was increased, ASDSF decreased. This was expected as with more data more nodes in the species tree become fully resolved, and the difference in support between CMCMC and UMCMC for those nodes will be zero. Conversely, ASDSF increased as the number of taxa was increased. Again this was expected, as denser taxon sampling will reduce the proportion of fully resolved nodes, and hence the proportion of nodes with zero difference between methods. In summary, while the underlying distributions sampled by CMCMC and UMCMC are different by construction, our results show that in practice they are almost the same.

If an internal node in one constraint gene tree (CGT) is binary, we consider such node as resolved. For young divergence time scenarios, there are fewer substitutions and hence less information available with which to reconstruct the phylogeny. For a given threshold, CGTs in young divergence time scenarios were less resolved than CGTs in old divergence time scenarios in Figure S5. But for all conditions the proportion of resolved nodes steadily decreased as the threshold was raised. A similar trend was observed when increasing the number of taxa in Figure S6. This shows the role that the support threshold plays as a useful tuning parameter for our heuristic.

While the ASDSF provides a numeric measure reflecting the similarity between the distributions being sampled, we visualize in Figure 4 the distributions sampled by the various samplers as an illustration of this similarity. While decreasing the support threshold increases the difference in the posterior distribution, all CMCMC methods approximate posterior density distributions similar to UMCMC, except for CMCMC-0.

**Figure 4:**
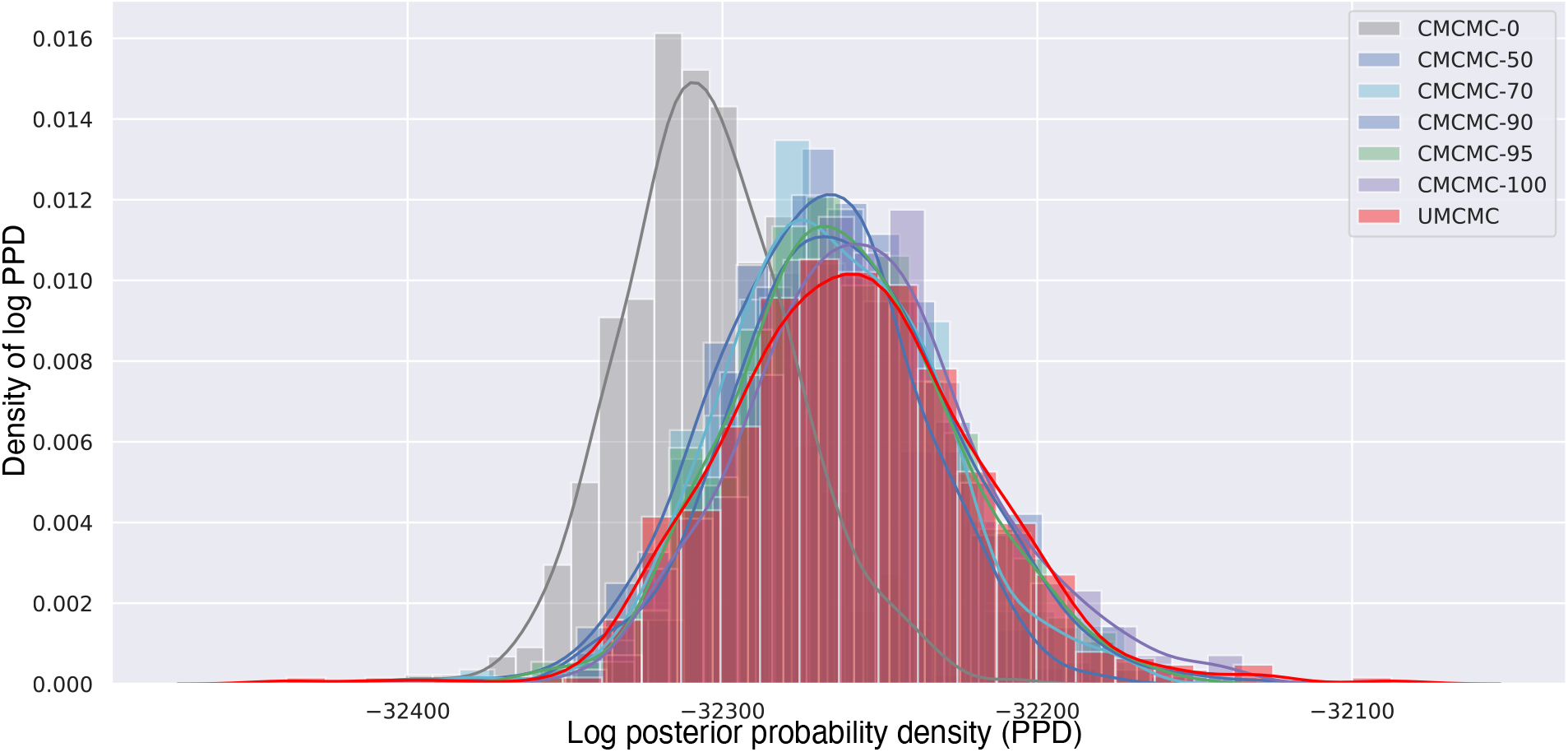
Kernel Density Estimation of the posterior probability distribution of CMCMC and UMCMC samples. Different methods are shown in different colors. All methods are run on the example sequence data which contains 16 taxa and 10 independent loci for the young divergence times and high population size scenario.

Our aim for CMCMC is to closely approximate the unconstrained posterior distribution of species trees, and to do so faster than UMCMC. The posterior density distribution of CMCMC-0 is very divergent from UMCMC and from CMCMC when using other thresholds, implying that it is no longer closely approximating the unconstrained posterior distribution. For this reason, we do not recommend using CMCMC-0 (i.e. gene trees with fixed, maximum likelihood estimated unrooted topologies) for Bayesian inference of species trees from sequences.

### 3.3 CMCMC and UMCMC predictions are essentially identical in accuracy

While Bayesian MCMC provides an approximation of the posterior distribution over parameters, the topology of the species tree is often the main quantity of interest. We compared the average Robinson-Foulds (RF) distance (Robinson and Foulds, 1981) between the true species tree and the inferred species tree topology in the 95% credible set to assess the topological accuracy. For each topology in the 95% credible set, we calculated the RF distance and divided it by the maximum possible RF distance (twice the number of internal branches in the species tree) to derive the normalized RF distance (normRF; Kupczok *et al*., 2010). Then we averaged all normRF distances, weighted by the frequency of each topology in the 95% credible set. More details about how to calculate average normRF distance are provided in the Evaluation Metrics section.

Figure 5 shows the topological accuracy of the species trees inferred by the various methods. As the figure shows, under both evolutionary scenarios, all samplers infer almost the same species trees. For the OH scenario the outlier corresponds to a single species tree which is difficult to accurately infer because of its short internal branches (Figure S7). The same lack of variation is also observed when varying the numbers of loci and taxa (Figure S8 and Figure S9).

**Figure 5:**
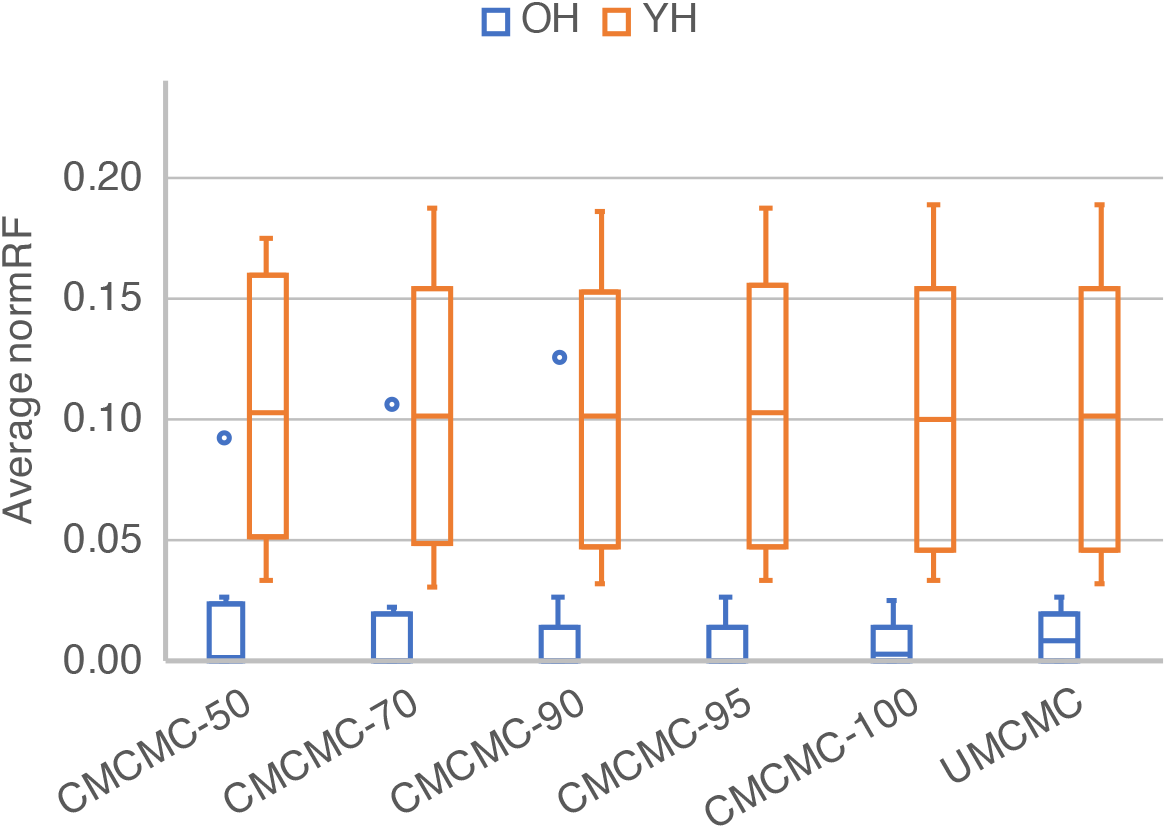
Topological accuracy of the species trees inferred by CMCMC and UMCMC. The data pertained to simulations of 10 loci from 16 taxa under the OH and YH scenarios.

Within a condition, there was no observed trend in average normRF distance (Figure 5, Figure S8 and Figure S9). In addition to topological error, we also calculated “branch length error” as the average branch score of trees in the credible set. The calculation is described in detail in the Evaluation Metrics subsection. As expected, branch length error was higher for the “old” case because branch score is not scale invariant, and average normRF distance was higher for the “young” case due to the lower rate of informative mutations. But within a condition, there was no difference in branch length error between CMCMC and UMCMC (Figure S10, Figure 11 and Figure S12). These results further demonstrate the proximity of the posterior distribution of species trees inferred by CMCMC to UMCMC.

### 3.4 Analysis of a biological data set

Recently, a study that applied exon capture sequencing to Australian rainbow skinks (Bragg *et al*., 2018) compared the phylogenies inferred using summary MSC methods, a full Bayesian MSC method and concatenation. However due to the computational time required, the full Bayesian species tree method was only applied to non-overlapping 32 locus subsets of the data, despite 304 highly informative loci being available. This data set contains 46 taxa from 43 recognized species.

CMCMC enabled us to double the number of loci, so inspired by Bragg *et al*., we compared the species trees inferred using UMCMC from nine non-overlapping 32 locus subsets with those inferred using CMCMC from four non-overlapping 64 locus subsets. All analyses were run until satisfactory convergence was observed. The samples for each analysis were summarized as maximum clade credibility (MCC) trees. To quantify the variation between species trees inferred from different subsets, we calculated the normRF distance between the MCC tree from one MCMC chain or the inferred tree from ASTRAL (Mirarab *et al*., 2014). As shown in Figure 6, CMCMC derived more consistent results compared with UMCMC, as the highest normRF distance between CMCMC subsets was 0.23, but the highest pairwise distance between the smaller UMCMC subsets was 0.3.

**Figure 6:**
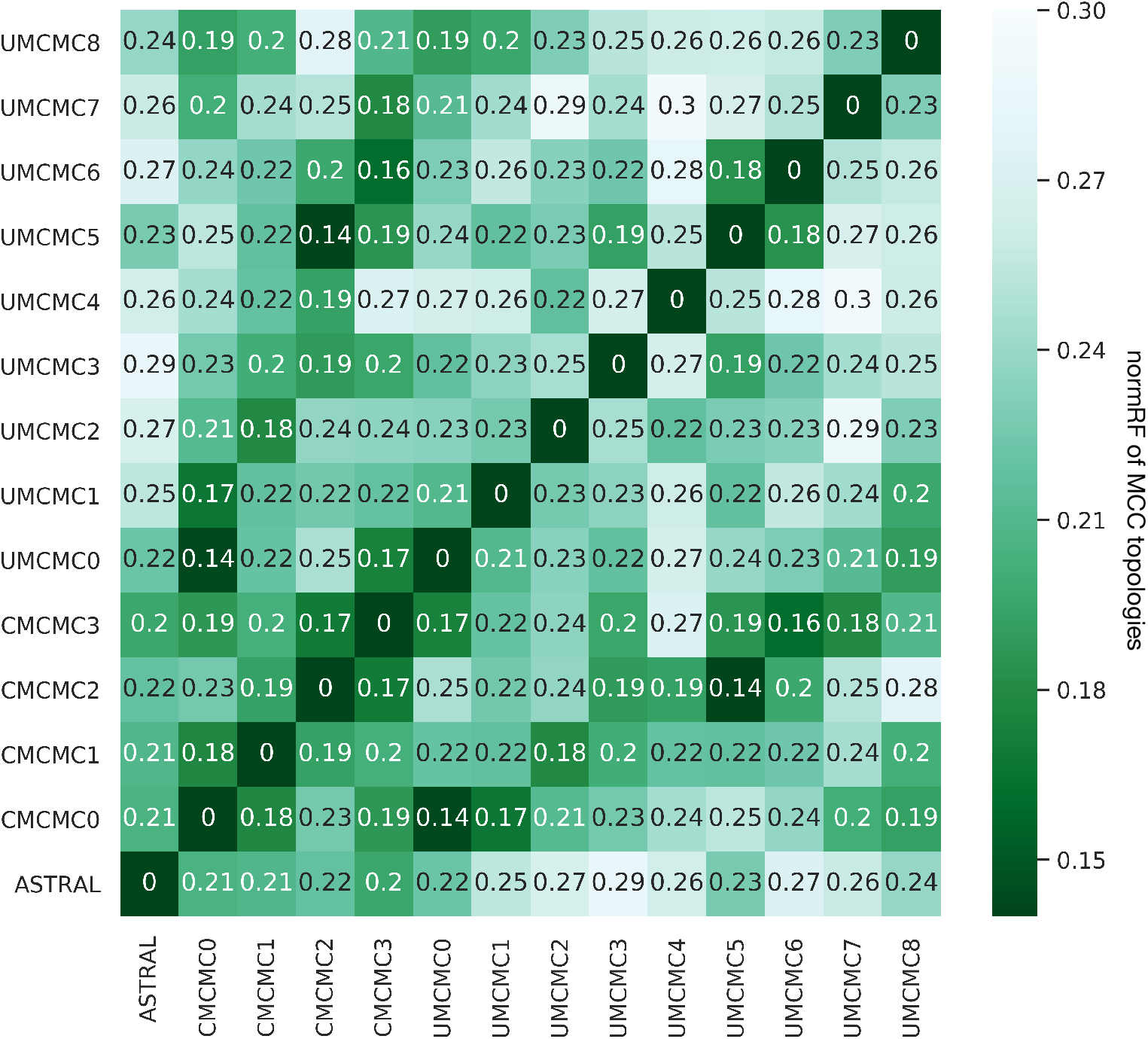
Discordance among phylogenies estimated by ASTRAL, CMCMC and UMCMC. CMCMC was applied to four non-overlapping subsets of 64 loci each, UMCMC was applied to nine non-overlapping subsets of 32 loci each, and ASTRAL was applied to gene trees inferred from all 304 loci. The color and the number in each entry of the matrix indicates the normalized Robinson-Foulds distance between maximum clade credibility (MCC) species trees estimated from each subset.

The precision of the 64 locus CMCMC posterior distributions was higher, as expected given the larger number of loci employed. For both normRF and branch score, the average distances between the maximum clade credibility (MCC) tree and individual samples in the posterior distribution were smaller for CMCMC (Figure S13 and Figure S14).

The increased precision of the CMCMC analyses enables taxonomic refinement of rainbow skinks. When 32 loci are used with UMCMC, the trio *Carlia inconnexa, C. pectoralis* and *C. rubigo* form a clade but the relationships within that clade are unclear, as the best supported topology for this trio has *C. rubigo* as sister with an average posterior probability of 69% across subsets. When 64 loci are used with CMCMC, this average rises to 98% (Figure S15 and Figure S16).

### 3.5 Conclusions

In this paper we reported on a simple heuristic method for speeding the convergence of Bayesian MCMC under the multispecies coalescent. The heuristic works by restricting the space of gene trees that can be sampled. The constraints can be obtained in various ways including bootstrap trees contracted according to some support threshold, majority-rule consensus trees of posterior samples, or even constraints provided based on biological knowledge. As the approach restricts the explored space by design, we evaluated the method’s performance in terms of convergence and, when converged, the distribution it samples. The evaluation was done on simulated data sets as well as a biological data set, and for evaluation metrics, we focused mainly on the time to convergence, the ASDSF between the constrained version of the sampler (CMCMC) and the unconstrained one (UMCMC), as well as the topological accuracy of the inferred species trees.

We have demonstrated that constraint gene trees are advantageous in two distinct ways. For datasets where it is challenging to achieve convergence with UMCMC, e.g. those which did not converge even after 20 days in our study, CMCMC can converge within a reasonable time. For datasets which did readily converge using UMCMC, applying constraint gene trees reduced the required number of iterations.

Following from our results, we recommend that CMCMC can be applied in two ways. The first is to accelerate preliminary analyses, where CMCMC-50 can be used to infer posterior distributions of species trees which are very close to the unconstrained posterior distributions. This has the additional benefit of reducing the amount of resources required for a given study, which range from grant money to pay for computer time, to CPU hours which may be in high demand at a given institution, to the electricity and natural resources needed to manufacture and operate computer nodes. As many preliminary analyses may have to be run for a given study, this reduction can be substantial.

The second is for final published analyses, where researches may wish to be more conservative and avoid approximations and heuristics wherever possible. For example, MCMC chains of finite lengths and variational Bayes are both approximate and heuristic methods, but are unavoidable for Bayesian inference of phylogenetic trees from sequences. Constraint gene trees are an additional heuristic which are avoidable for datasets where convergence is possible within a reasonable time using UMCMC, but unavoidable otherwise. So our final recommendation is to employ CMCMC for final published analyses of datasets which fail to converge using UMCMC.

## 4 Materials and Methods

### 4.1 MCMC implementation and settings

CMCMC can be easily implemented in commonly used Bayesian based phylogenetics software packages such as PhyloNet (Wen *et al*., 2016), BEAST (Drummond and Rambaut, 2007) and BEAST2.5 (Bouckaert *et al*., 2019). In this paper, we implemented CMCMC in PhyloNet. In all our experiments, we generated 50 bootstrap trees for each locus and obtained the majority-rule consensus trees from those. Firstly, we generated bootstrap trees given an alignment using RAxML (Stamatakis, 2014). Then, we estimated the constraint tree given a specific support threshold.

We applied a uniform prior over the species tree topologies, a uniform prior *U*(0, ∞) on species tree node heights, and a 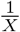 prior on the mean population size. More implementation details are provided in Supplementary Materials.

### 4.2 Executing MCMC chains

For the simulation study, we first ran each chain for three days on the DAVinCI computing cluster. All jobs executed on this cluster ran on 2.83GHz Intel Westmere CPUs. Jobs were restarted each day, and so the total running time of the MCMC chain was less than 72 hours, as for each job some time was spent queuing and postprocessing.

For any chain that did not converge within three days, we restarted it from the beginning on the NOTS computing cluster at Rice University. Jobs executed on this cluster were randomly assigned to one of the following CPUs: Intel Xeon E5-2650 v2 at 2.6GHz, Xeon E5-2650 v4 at 2.2GHz, Xeon Gold 6126 at 2.6GHz, or Intel Xeon Gold 6230 at 2.1GHz. With the exception of the 48 taxon analyses, all chains were run for 20 days. The 48 taxon chains were run for only 10 days because we noticed they had all converged by that time. As with the shorter chains, jobs were restarted each day so the total runtime was less than 480 hours, and less than 240 hours for the 48 taxon chains.

For the empirical study, 10 independent chains with different random seeds but otherwise identical data and settings were run for each locus subset and method. This was necessary to achieve convergence on these relatively large datasets. CMCMC chains on 64 loci were run for 160 million iterations, taking approximately 25 days. UMCMC chains on 32 loci were run for 120 million iterations, taking approximately 10 days. After all 10 chains had finished running for a given subset and method, the remaining samples were concatenated after removing the 10% burnin.

### 4.3 Simulating data

For all simulated data sets, we used DendroPy (Sukumaran and Holder, 2010) to generate random species trees and ms (Hudson, 2002) to generate gene trees on these species trees under the multispecies coalescent. Sequence data were generated by Seq-Gen (Rambaut and Grass, 1997) under the Jukes-Cantor model(Jukes and Cantor, 1969). We derived the CGT for each locus by bootstrapping from the sequences by RAxML (Stamatakis, 2014).

Because species tree inference methods are employed over a range of evolutionary time scales and to clades with different population sizes, we varied both parameters for our simulation study which are shown in Table 1. For each simulated species tree we scaled its root to different heights; an “old” height of 50 million years ago (mya) akin to the rice–Pooideae split (Sandve *et al*., 2008), and a “young” height of 10 mya akin to the split of gorillas with humans and chimpanzees (Langergraber *et al*., 2012). Gene trees were scaled before simulating sequences so that the branch lenghts in substitutions per site corresponded to a substitution rate of 2.5 × 10^−3^ per million years. This rate is slightly faster than the rates observed for the RAG1 nuclear gene in animals (Hugall *et al*., 2007), but within the ranges observed for plant nuclear genes (Huang *et al*., 2003).

**Table 1:** The evolutionary parameters varied to control the complexity and signal in the data.

|  | Low population sizes | High population sizes |
| --- | --- | --- |
| Old divergence times | OL: 50 mya, 100k individuals | OH: 50 mya, 500k individuals |
| Young divergence times | YL: 10 mya, 100k individuals | YH: 10 mya, 500k individuals |

For both the old and young species tree scales, we simulated gene trees under large and small population sizes of 500,000 and 100,000 respectively, with annual generation times. Chimpanzees, gorillas and ancient humans all have effective population sizes (*N_e_*) of around 20,000 individuals (Huff *et al*., 2010). Assuming a great ape generation time of 25 years, this population will have the same distribution of coalescent times as a clade of species with annual generation times and an *N_e_* of 500,000, the same as our large population size condition. Given that human effective population sizes are often considered low, the small population size condition therefore corresponds to species with very low effective population sizes.

The evolutionary parameters also affect the proportion of resolved internal nodes of the CGT as shown in Figure S5. The “OH” and “OL” scenarios have higher proportion of resolved internal nodes than the “YH” and “YL” which means that the substitution rate more effectively restricts the gene tree search space. The proportion of resolved internal nodes in consensus trees decreases as the number of taxa increases as shown in Figure S6. In contrast, the population size does not have such obvious effect as the substitution rate or number of taxa.

More details on the simulations are provided in Supplementary Materials.

### 4.4 Biological data

We analyzed the Australian skinks data set which is provided in Bragg *et al*. 2018. We randomly selected one sample from each species. Note that the species names in the data set and in the paper are not consistent. More details about how to map the species in the data set and in the paper are shown in Table S1.

The Australian skinks data set contains three in-group genera: *Carlia, Lygisaurus* and *Liburnascincus*. There are 46 taxa from 43 recognized species. All details of the biological data including genus, species, tissue, collection, sample library and focal clade are provided in supplementary Table S2.

To obtain informative gene trees, we included 304 complete informative loci whose length ranges from 240 to 6,534 sites. Figure S17 shows the proportion of resolved internal nodes of constraint gene trees for different ranges of sequence length. In general, as the length of sequence increases the number of resolved internal nodes gets larger. This is because longer sequences are likely to contain more substitutions to inform the resolution of nodes.

### 4.5 Evaluation metrics

#### 4.5.1 Effective sample size

The Effective Sample Size or ESS is the number of effectively independent draws from some distributions sampled by the MCMC chain. Adequate ESS is a sign of good mixing of the MCMC chain and it has been argued that the ESS should be more than 200 (Kuhner, 2009), a value that has been adopted in the Bayesian phylogenetics community. Therefore, an MCMC chain where the ESS of all selected probability densities and parameter values were higher than 200 was considered to have converged. The probability densities were of the posterior, phylogenetic likelihood, prior and coalescent likelihood, all of which are dependent on the tree topologies and continuous parameters. The parameter values were tree height and population size.

#### 4.5.2 Average Robinson-Foulds distance

When calculating the total Robinson-Foulds distance, we only considered posterior samples where the species tree topology was within the 95% credible set. We call this credible set of posterior samples *T*\* to distinguish it from the full set of posterior samples *T*. To quantify differences between true tree *t* and the 95% credible set *T*\*, we calculate the average normRF distance (Kupczok *et al*., 2010) as

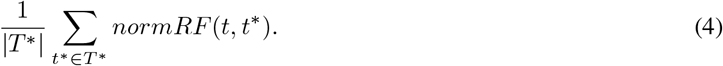

#### 4.5.3 Average branch length error

To evaluate the accuracy of branch length estimates, we calculated the average error between the true tree t and the 95% credible set *T*\* using a measure based on Euclidean branch score distances (Kuhner and Felsenstein, 1994; St. John, 2017). For every tree *t*\* in the credible set, we take the union *B* of all branches in *t*\* and *t*, where a branch *b* ∈ *B* is defined by the taxa associated with its tipward node (i.e. the corresponding clade). We define *δ*(*b*) as the difference between the length of *b* in *t* and *t*\*. If a branch is missing in one of *t* and *t*\*, its length in that tree is defined to be zero. The sum of square differences 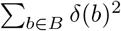 is known as the branch score distance, which we will treat as a function *BSD*(*t*, *t*\*). The square root of the branch score is a Euclidean distance, and we define branch length error as

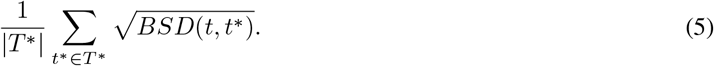

#### 4.5.4 Average standard deviation of split frequencies

Average standard deviation of split frequencies (ASDSF) is a measure of convergence that has been used in tools such as ExaBayes (Aberer *et al*., 2014) and MrBayes (Ronquist *et al*., 2012). ASDSF can be calculated by comparing split or clades frequencies between two MCMC chains. Given two posterior distributions *T*_1_ and *T*_2_ from two MCMC chains and their corresponding 95% credible sets 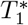 and 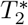, *C* is all unique, non-trivial clades in 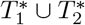. Set *C*\* is defined as

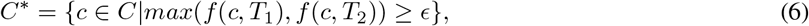

where *f*(*c,T*) is the frequency of clade *c* in distribution *T*, and *ϵ* is a threshold. We used *ϵ* = 0.1, the same as the default setting in MrBayes 3.2 (Ronquist *et al*., 2012). Finally, the ASDSF between *T*_1_ and *T*_2_ is defined as

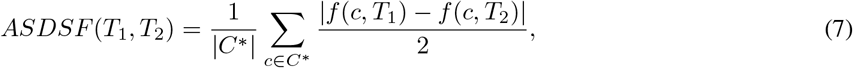

because the standard deviation of two numbers is half of the absolute difference.

## 5 Supplementary Material

Supplementary Figures S1-S18, Tables S1-S2 and external tool commands are available in supplementary.pdf. All simulation data are available online: https://drive.google.com/file/d/1T56Hz3tMCkMU0qXs8-CftFJwsooKwBao/view?usp=sharing.

## 6 Acknowledgments

This work was supported in part by NSF grants DBI-1355998, CCF-1514177, CCF-1800723, DMS-1547433, and by the Data Analysis and Visualization Cyberinfrastructure funded by NSF under grant OCI-0959097 and Rice University. Ana C. Afonso Silva assisted us in interpreting and mapping the species names of the biological data set.

## References

Aberer, A. J., Kobert, K., and Stamatakis, A. 2014. ExaBayes: massively parallel Bayesian tree inference for the whole-genome era. Molecular Biology and Evolution, 31(10): 2553–2556.

Attias, H. 1999. Inferring parameters and structure of latent variable models by variational Bayes. In Proceedings of the Fifteenth Conference on Uncertainty in Artificial Intelligence, UAI’99, pages 21–30, San Francisco, CA, USA. Morgan Kaufmann Publishers Inc.

Bouckaert, R., Vaughan, T. G., Barido-Sottani, J., Duchêne, S., Fourment, M., Gavryushkina, A., Heled, J., Jones, G., Kühnert, D., De Maio, N., Matschiner, M., Mendes, F. K., Müller, N. F., Ogilvie, H. A., du Plessis, L., Popinga, A., Rambaut, A., Rasmussen, D., Siveroni, I., Suchard, M. A., Wu, C.-H., Xie, D., Zhang, C., Stadler, T., and Drummond, A. J. 2019. BEAST 2.5: An advanced software platform for Bayesian evolutionary analysis. PLOS Computational Biology, 15(4): 1–28.

Bragg, J. G., Potter, S., Silva, A. C. A., Hoskin, C. J., Bai, B. Y., and Moritz, C. 2018. Phylogenomics of a rapid radiation: the Australian rainbow skinks. BMC Evolutionary Biology, 18(1): 15.

Chifman, J. and Kubatko, L. 2014. Quartet inference from SNP data under the coalescent model. Bioinformatics, 30(23): 3317–3324.

Degnan, J. H. and Rosenberg, N. A. 2009. Gene tree discordance, phylogenetic inference and the multispecies coalescent. Trends in Ecology & Evolution, 24(6): 332–340.

Drummond, A. J. and Rambaut, A. 2007. BEAST: Bayesian evolutionary analysis by sampling trees. BMC Evolutionary Biology, 7(1): 214.

Felsenstein, J. 1978. The number of evolutionary trees. Systematic Biology, 27(1): 27–33.

Flouri, T., Jiao, X., Rannala, B., and Yang, Z. 2018. Species tree inference with BPP using genomic sequences and the multispecies coalescent. Molecular Biology and Evolution, 35(10): 2585–2593.

Fourment, M. and Darling, A. E. 2019. Evaluating probabilistic programming and fast variational Bayesian inference in phylogenetics. bioRxiv.

Geyer, C. J. 1991. Markov Chain Monte Carlo Maximum Likelihood. In E. M. Keramidas, editor, Computing Science and Statistics: Proceedings of the 23rd Symposium on the Interface, pages 156–163.

Höhna, S. and Drummond, A. J. 2011. Guided Tree Topology Proposals for Bayesian Phylogenetic Inference. Systematic Biology, 61(1): 1–11.

Huang, S., Su, X., Haselkorn, R., and Gornicki, P. 2003. Evolution of switchgrass (Panicum virgatum L.) based on sequences of the nuclear gene encoding plastid acetyl-CoA carboxylase. Plant Science, 164(1): 43–49.

Hudson, R. R. 2002. Generating samples under a Wright–Fisher neutral model of genetic variation. Bioinformatics, 18(2): 337–338.

Huelsenbeck, J. P., Ronquist, F., Nielsen, R., and Bollback, J. P. 2001. Bayesian inference of phylogeny and its impact on evolutionary biology. Science, 294(5550): 2310–2314.

Huff, C. D., Xing, J., Rogers, A. R., Witherspoon, D., and Jorde, L. B. 2010. Mobile elements reveal small population size in the ancient ancestors of Homo sapiens. Proceedings of the National Academy of Sciences, 107(5): 2147–2152.

Hugall, A. F., Foster, R., and Lee, M. S. Y. 2007. Calibration Choice, Rate Smoothing, and the Pattern of Tetrapod Diversification According to the Long Nuclear Gene RAG-1. Systematic Biology, 56(4): 543–563.

Jones, G. 2017. Algorithmic improvements to species delimitation and phylogeny estimation under the multispecies coalescent. Journal of Mathematical Biology, 74(1): 447–467.

Jukes, T. H. and Cantor, C. R. 1969. Evolution of Protein Molecules. In Mammalian Protein Metabolism, pages 21–132. Academic Press.

Kuhner, M. K. 2009. Coalescent genealogy samplers: windows into population history. Trends in Ecology & Evolution, 24(2): 86–93.

Kuhner, M. K. and Felsenstein, J. 1994. A simulation comparison of phylogeny algorithms under equal and unequal evolutionary rates. Molecular Biology and Evolution, 11(3): 459–468.

Kupczok, A., Schmidt, H. A., and von Haeseler, A. 2010. Accuracy of phylogeny reconstruction methods combining overlapping gene data sets. Algorithms for Molecular Biology, 5(1): 37.

Lakner, C., Van Der Mark, P., Huelsenbeck, J. P., Larget, B., and Ronquist, F. 2008. Efficiency of Markov chain Monte Carlo tree proposals in Bayesian phylogenetics. Systematic Biology, 57(1): 86–103.

Langergraber, K. E., Prüfer, K., Rowney, C., Boesch, C., Crockford, C., Fawcett, K., Inoue, E., Inoue-Muruyama, M., Mitani, J. C., Muller, M. N., Robbins, M. M., Schubert, G., Stoinski, T. S., Viola, B., Watts, D., Wittig, R. M., Wrangham, R. W., Zuberbühler, K., Pääbo, S., and Vigilant, L. 2012. Generation times in wild chimpanzees and gorillas suggest earlier divergence times in great ape and human evolution. Proceedings of the National Academy of Sciences, 109(39): 15716–15721.

Liu, L. and Yu, L. 2011. Estimating species trees from unrooted gene trees. Systematic Biology, 60(5): 661–667.

Liu, L., Yu, L., and Edwards, S. V. 2010. A maximum pseudo-likelihood approach for estimating species trees under the coalescent model. BMC Evolutionary Biology, 10(1): 302.

Mazza, G., Menchetti, M., Sluys, R., Sola, E., Riutort, M., Tricarico, E., Justine, J.-L., Cavigioli, L., and Mori, E. 2016. First report of the land planarian Diversibipalium multilineatum (Makino & Shirasawa, 1983)(Platyhelminthes, Tricladida, Continenticola) in Europe. Zootaxa, 4067(5): 577–580.

Mirarab, S., Reaz, R., Bayzid, M. S., Zimmermann, T., Swenson, M. S., and Warnow, T. 2014. ASTRAL: genome-scale coalescent-based species tree estimation. Bioinformatics, 30(17): i541–i548.

Ogilvie, H. A., Heled, J., Xie, D., and Drummond, A. J. 2016. Computational performance and statistical accuracy of *BEAST and comparisons with other methods. Systematic Biology, 65(3): 381–396.

Ogilvie, H. A., Bouckaert, R. R., and Drummond, A. J. 2017. StarBEAST2 brings faster species tree inference and accurate estimates of substitution rates. Molecular Biology and Evolution, 34(8): 2101–2114.

Rambaut, A. and Grass, N. C. 1997. Seq-Gen: an application for the Monte Carlo simulation of DNA sequence evolution along phylogenetic trees. Bioinformatics, 13(3): 235–238.

Rambaut, A., Drummond, A. J., Xie, D., Baele, G., and Suchard, M. A. 2018. Posterior summarization in Bayesian phylogenetics using Tracer 1.7. Systematic Biology, 67(5): 901–904.

Rannala, B. and Yang, Z. 2017. Efficient Bayesian Species Tree Inference under the Multispecies Coalescent. Systematic Biology, 66(5): 823–842.

Robert, C. and Casella, G. 2011. A Short History of Markov Chain Monte Carlo: Subjective Recollections from Incomplete Data. Statistical Science, 26(1): 102–115.

Robinson, D. F. and Foulds, L. R. 1981. Comparison of phylogenetic trees. Mathematical Biosciences, 53(1-2): 131–147.

Ronquist, F., Teslenko, M., van der Mark, P., Ayres, D. L., Darling, A., Höhna, S., Larget, B., Liu, L., Suchard, M. A., and Huelsenbeck, J. P. 2012. MrBayes 3.2: efficient Bayesian phylogenetic inference and model choice across a large model space. Systematic Biology, 61(3): 539–542.

Sandve, S. R., Rudi, H., Asp, T., and Rognli, O. A. 2008. Tracking the evolution of a cold stress associated gene family in cold tolerant grasses. BMC Evolutionary Biology, 8(1): 245.

St. John, K. 2017. Review Paper: The Shape of Phylogenetic Treespace. Systematic Biology, 66(1): e83–e94.

Stamatakis, A. 2014. RAxML version 8: a tool for phylogenetic analysis and post-analysis of large phylogenies. Bioinformatics, 30(9): 1312–1313.

Stensvold, C. R., Lebbad, M., and Clark, C. G. 2011. Last of the human protists: the phylogeny and genetic diversity of Iodamoeba. Molecular Biology and Evolution, 29(1): 39–42.

Stunženas, V., Petkevičiūtė, R., and Stanevižiūtė, G. 2011. Phylogeny of *Sphaerium solidum* (Bivalvia) based on karyotype and sequences of 16S and ITS1 rDNA. Central European Journal of Biology, 6(1): 105–117.

Sukumaran, J. and Holder, M. T. 2010. DendroPy: a Python library for phylogenetic computing. Bioinformatics, 26(12): 1569–1571.

Tajima, F. 1983. Evolutionary relationship of DNA sequences in finite populations. Genetics, 105(2): 437–460.

Takahata, N., Satta, Y., and Klein, J. 1995. Divergence time and population size in the lineage leading to modern humans. Theoretical Population Biology, 48(2): 198–221.

Wang, Y. and Nakhleh, L. K. 2018. Towards an accurate and efficient heuristic for species/gene tree co-estimation. Bioinformatics, 34 17: i697–i705.

Wen, D. and Nakhleh, L. 2017. Coestimating reticulate phylogenies and gene trees from multilocus sequence data. Systematic Biology, 67(3): 439–457.

Wen, D., Yu, Y., and Nakhleh, L. 2016. Bayesian inference of reticulate phylogenies under the multispecies network coalescent. PLOS Genetics, 12(5): e1006006.

Wen, D., Yu, Y., Zhu, J., and Nakhleh, L. 2018. Inferring phylogenetic networks using PhyloNet. Systematic Biology, 67(4): 735–740.

Yang, Z. 2002. Likelihood and Bayes estimation of ancestral population sizes in hominoids using data from multiple loci. Genetics, 162(4): 1811–1823.

Yang, Z. and Rannala, B. 2014. Unguided species delimitation using DNA sequence data from multiple loci. Molecular Biology and Evolution, 31(12): 3125–3135.

Yang, Z. and Rodríguez, C. E. 2013. Searching for efficient Markov chain Monte Carlo proposal kernels. Proceedings of the National Academy of Sciences, 110(48): 19307–19312.

Yu, Y., Dong, J., Liu, K. J., and Nakhleh, L. 2014. Maximum likelihood inference of reticulate evolutionary histories. Proceedings of the National Academy of Sciences, 111(46): 16448–16453.

Zhang, C. and Matsen, F. A. 2019. Variational Bayesian phylogenetic inference. In International Conference on Learning Representations. https://openreview.net/forum?id=SJVmjjR9FX accessed September 15 2019.

Zhang, C., Ogilvie, H. A., Drummond, A. J., and Stadler, T. 2017. Bayesian inference of species networks from multilocus sequence data. Molecular Biology and Evolution, 35(2): 504–517.

Zhang, C., Huelsenbeck, J. P., and Ronquist, F. 2019. Using parsimony-guided tree proposals to accelerate convergence in Bayesian phylogenetic inference. bioRxiv.

Zhu, J., Wen, D., Yu, Y., Meudt, H. M., and Nakhleh, L. 2018. Bayesian inference of phylogenetic networks from bi-allelic genetic markers. PLOS Computational Biology, 14(1): e1005932.

