## Supplementary information for "Practical Speedup of Bayesian Inference of Species Phylogenies by Restricting the Space of Gene Trees"

### S1 Supplementary Figures

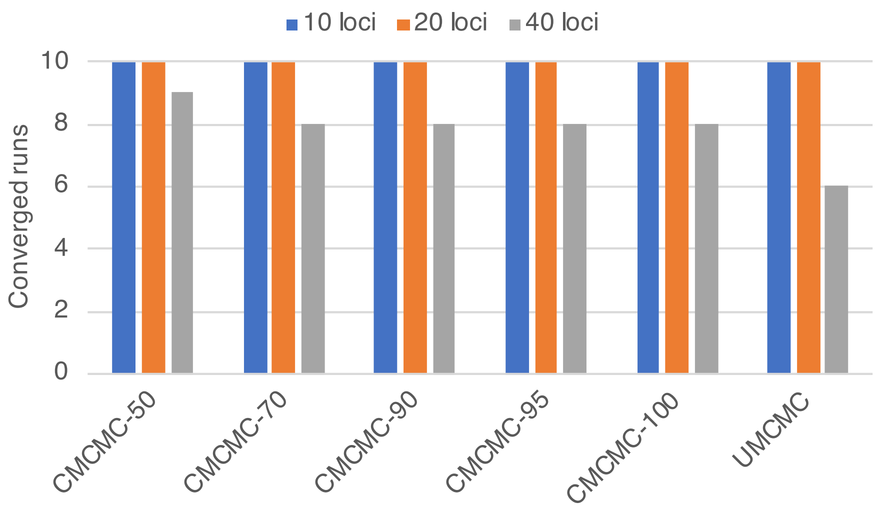


Figure S1: **Convergence of different methods for different numbers of loci, restricted to the "YH" scenario.** Different samplers are shown on the x axis and the y axis shows the number of replicates (out of 10) on which the sampler converged within 72 hours for 10 and 20 loci, and 20 days for 40 loci.

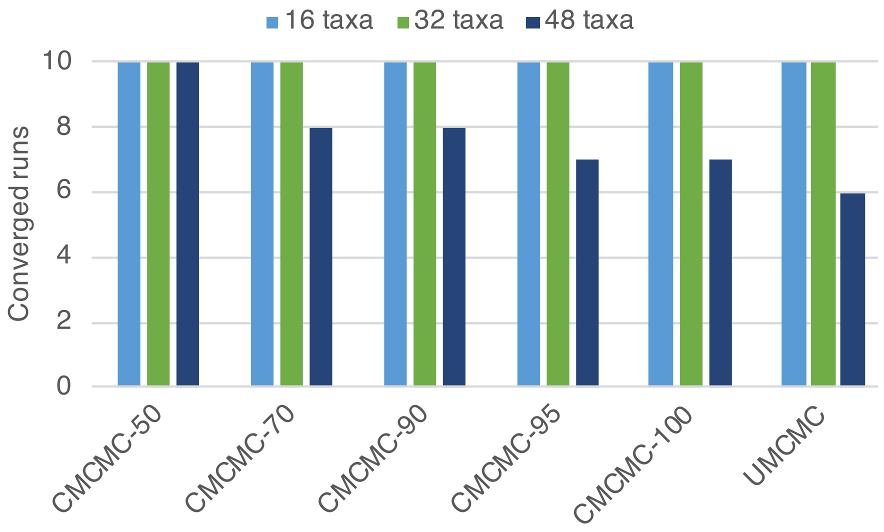


Figure S2: **Convergence of different methods for different numbers of taxa in the true species tree, restricted to the "YH" scenario.** Different samplers are shown on the x axis and the y axis shows the number of replicates (out of 10) on which the sampler converged within 72 hours

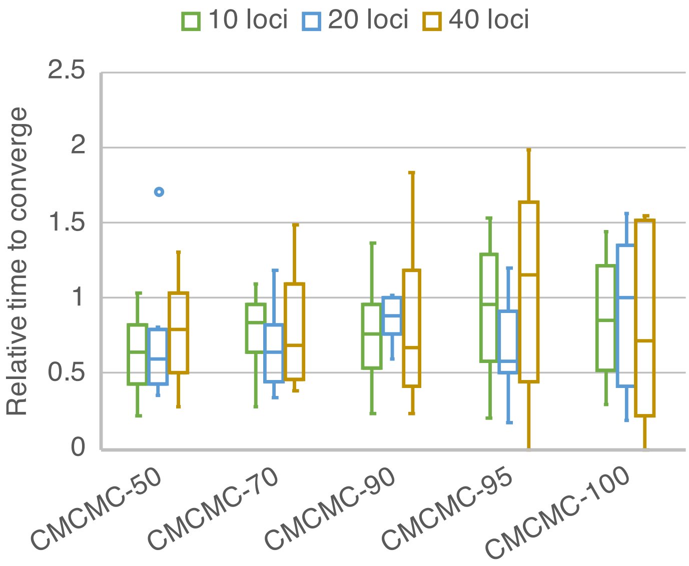


Figure S3: **Computational cost of different methods for different numbers of loci, restricted to the "YH" scenario.** The computational cost is shown as the ratios of iterations required for convergence using CMCMC compared with UMCMC. Values above 1 are replicates where UMCMC is faster than CMCMC, below 1 are replicates where CMCMC is faster than UMCMC.

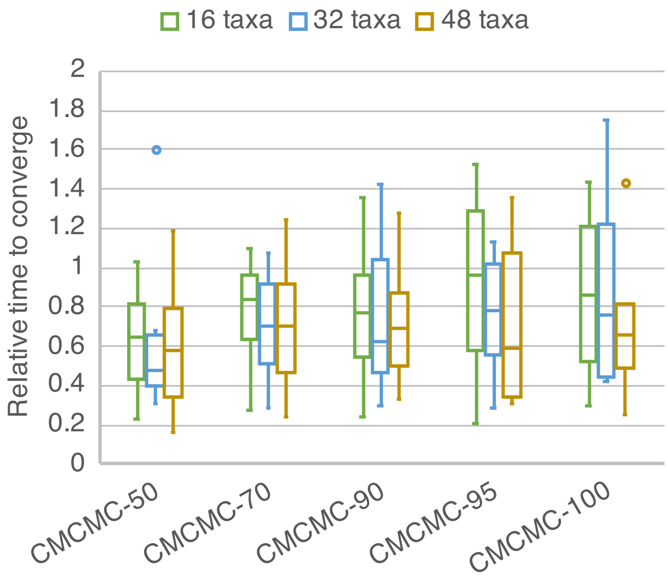


Figure S4: **Computational cost of different methods for different numbers of taxa in the species tree, restricted to the "YH" scenario.** The computational cost is shown as the ratios of iterations required for convergence using CMCMC compared with UMCMC. Values above 1 are replicates where UMCMC is faster than CMCMC, below 1 are replicates where CMCMC is faster than UMCMC.

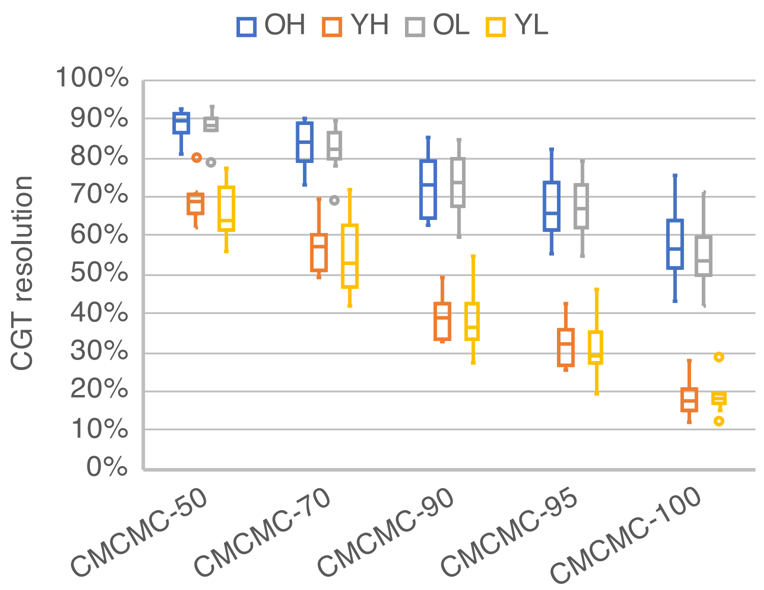


Figure S5: **The average proportion of resolved internal nodes for constraint gene trees (CGT) for replicates of different conditions.** Different colors represent different evolutionary conditions according to Table 1. There are 10 replicates for each condition. For each replicate, the resolution was averaged over 200 independent loci.

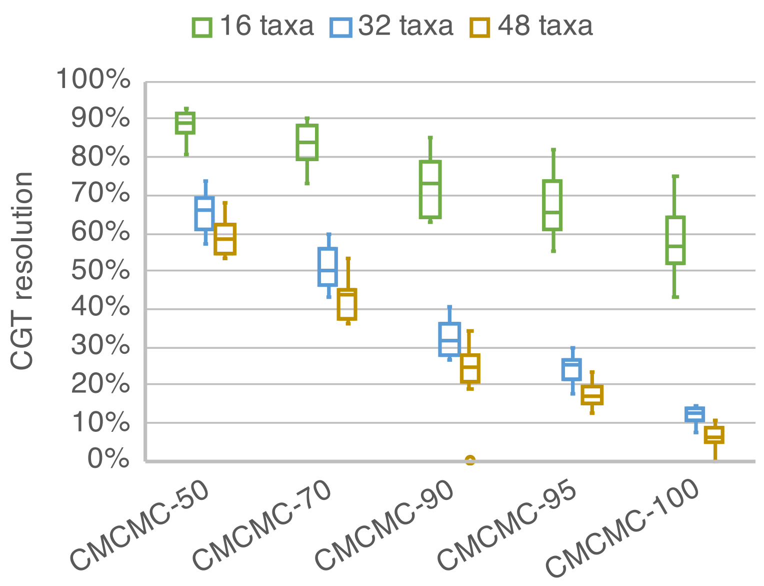


Figure S6: **The average proportion of resolved internal nodes for constraint gene trees (CGT) for replicates of different numbers of taxa, restricted to the "YH" scenario.** Different colors represent different numbers of taxa in the true species tree. The numbers of taxa are 16, 32 and 48. There are 10 replicates for each number of taxa. For each replicate, the resolution was averaged over 200 independent loci.

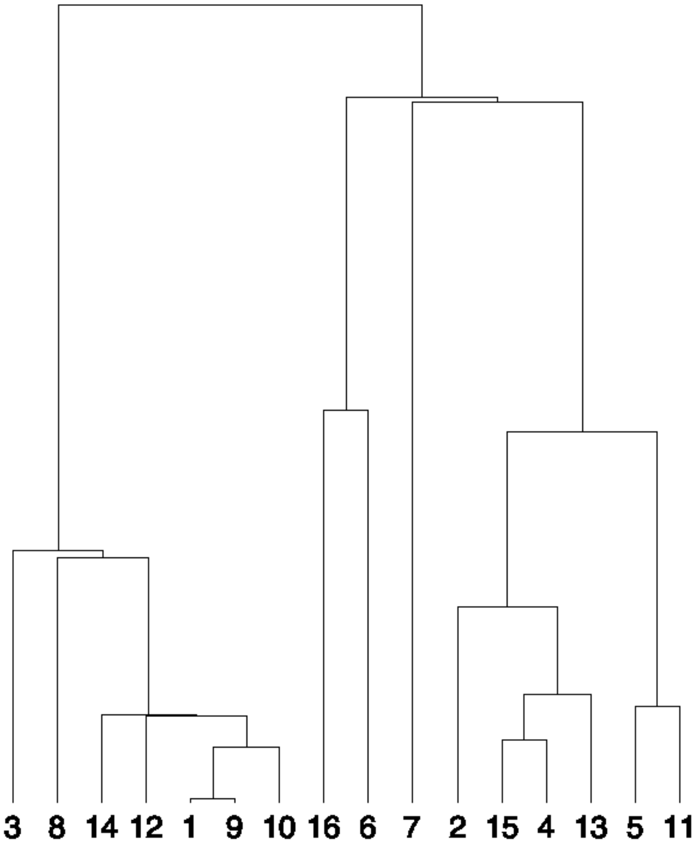


Figure S7: **The true species tree of the outlier of "OH" condition in Figure 5.**

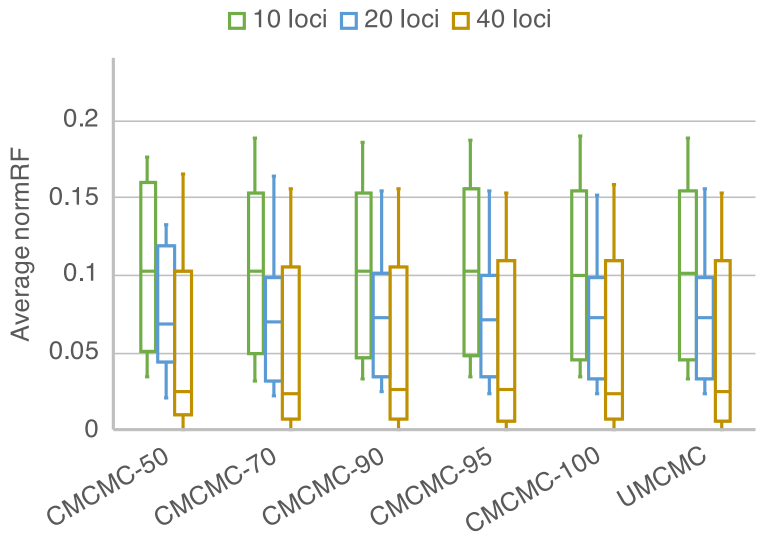


Figure S8: **Topological accuracy of CMCMC and UMCMC for different numbers of gene loci.** The x axis lists CMCMC samplers with different consensus thresholds and UMCMC. The y axis shows the averaged Robinson-Foulds (RF) distance. This figure shows the accuracy for 16-taxon species tree under "young divergence times" and "high population size".

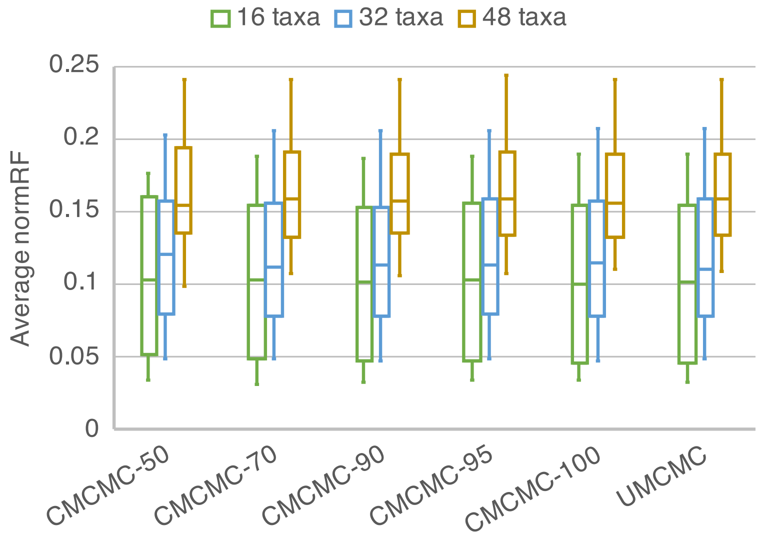


Figure S9: **Topological accuracy of CMCMC and UMCMC for different numbers of taxa in the true species tree.** The x axis lists CMCMC samplers with different consensus thresholds and UMCMC. The y axis shows the averaged Robinson-Foulds (RF) distance. This figure shows the accuracy for 10 independent loci under "young divergence times" and "high population size".

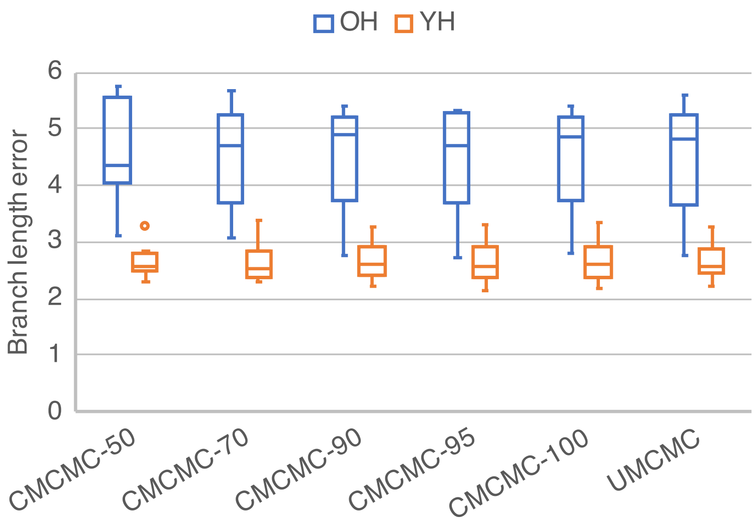


Figure S10: **Branch length error of CMCMC and UMCMC for different evolutionary scenarios.** The x axis lists CMCMC samplers with different consensus thresholds and UMCMC. The y axis shows the branch length error in coalescent units. This figure shows the branch length estimation when varying the divergence times fixing the number of taxa and loci as 16 and 10. Only ‘YH’ and ‘OH’ are shown because UMCMC cannot converge in ‘YL’ and ‘OL’ scenarios.

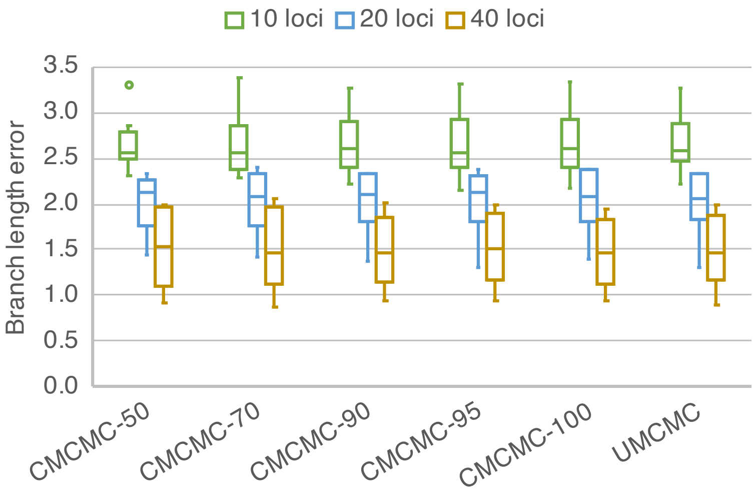


Figure S11: **Branch length error of CMCMC and UMCMC for different numbers of gene loci.** The x axis lists CMCMC samplers with different consensus thresholds and UMCMC. The y axis shows the branch length error in coalescent units. This figure shows the branch length estimation for 16-taxon species tree with 10, 20 and 40 loci under “young divergence times” and “high population size”.

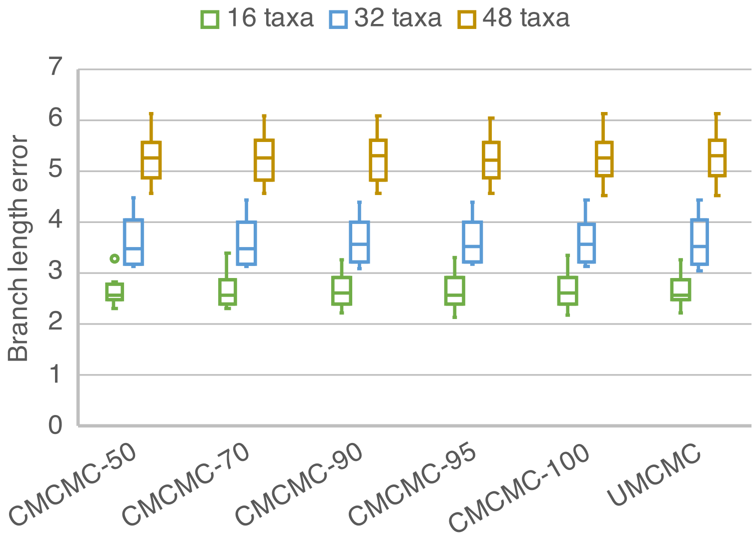


Figure S12: **Branch length error of CMCMC and UMCMC for different numbers of taxa in the true species tree.** The x axis lists CMCMC samplers with different consensus thresholds and UMCMC. The y axis shows the branch length error in coalescent units. This figure shows the branch length accuracy for 10 independent loci under "young divergence times" and "high population size".

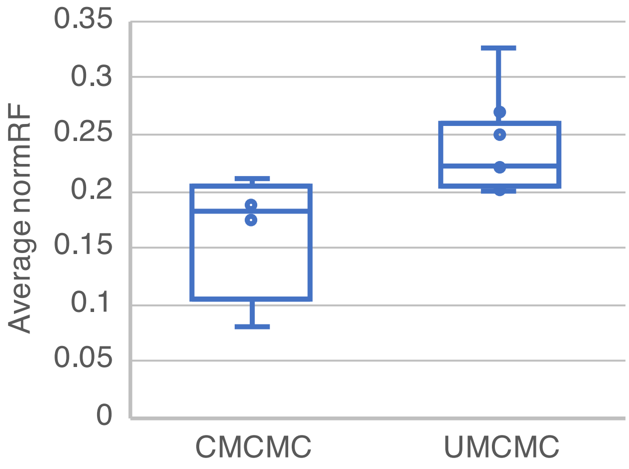


Figure S13: **Species tree topological precision of CMCMC and UMCMC on biological data.** The y axis shows the average normRF distance between the maximum clade credibility summary tree of all chains for a given method, and each individual sample for the corresponding method. CMCMC contains 4 chains and 225 samples are selected from each chain. UMCMC contains 9 chains and 100 samples are selected from each chain. For each method, there are 900 samples in total.

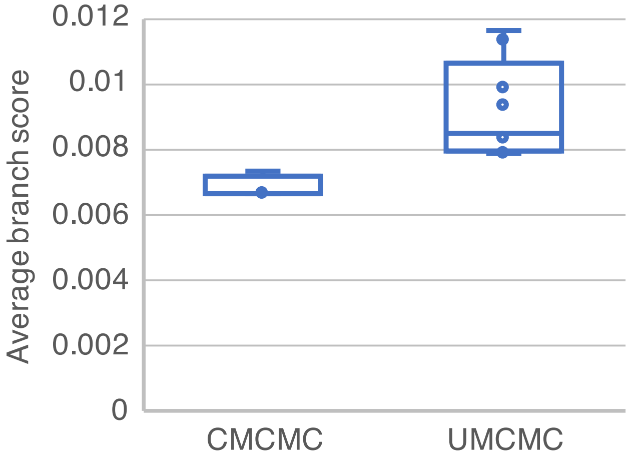


Figure S14: **Species tree branch length precision of CMCMC and UMCMC on biological data.** The y axis shows the average Euclidean branch score (Kuhner and Felsenstein, 1994; St. John, 2017) between the maximum clade credibility summary tree of all chains for a given method, and each individual sample for the corresponding method. The units are in substitutions per site. CMCMC contains 4 chains and 225 samples are selected from each chain. UMCMC contains 9 chains and 100 samples are selected from each chain. For each method, there are 900 samples in total.

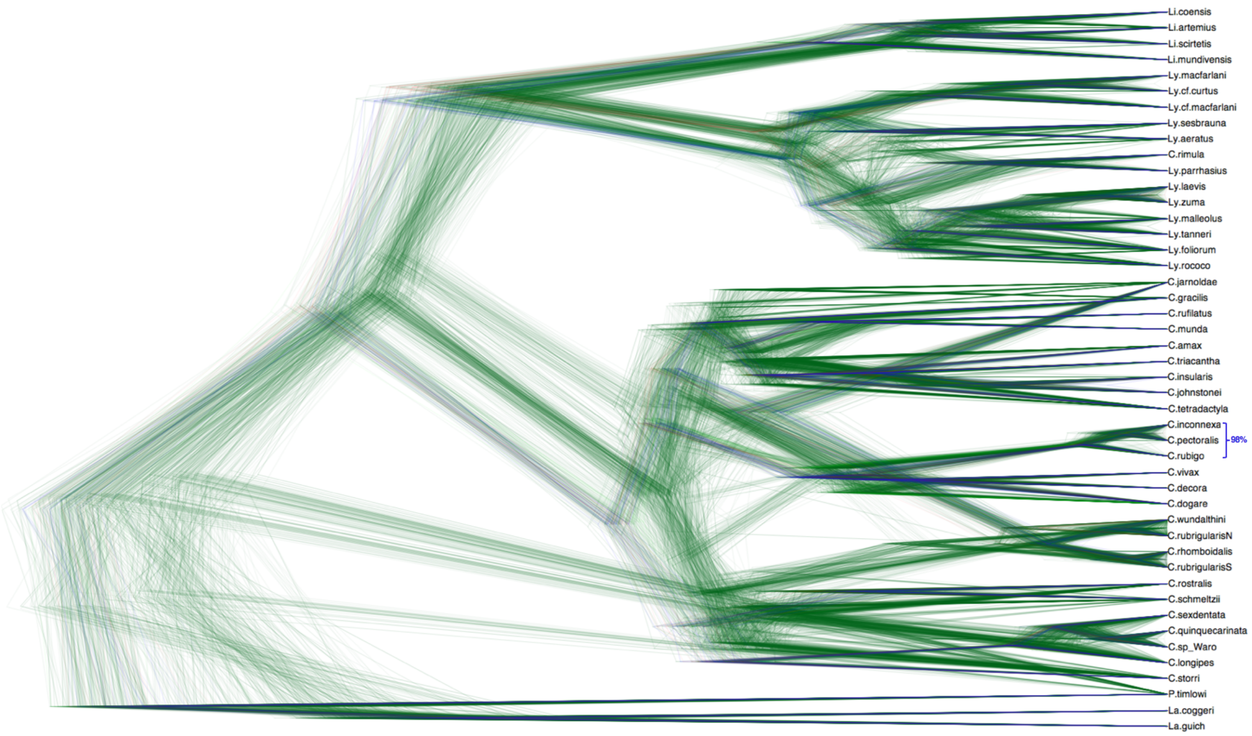


Figure S15: **Cloudogram of species tree samples from all subsets using CMCMC.** There are 900 samples shown in the above figure. For CMCMC, the 304 loci were separated into 4 subsets and 225 samples per subset were randomly selected from each subset. The consensus threshold is 50.

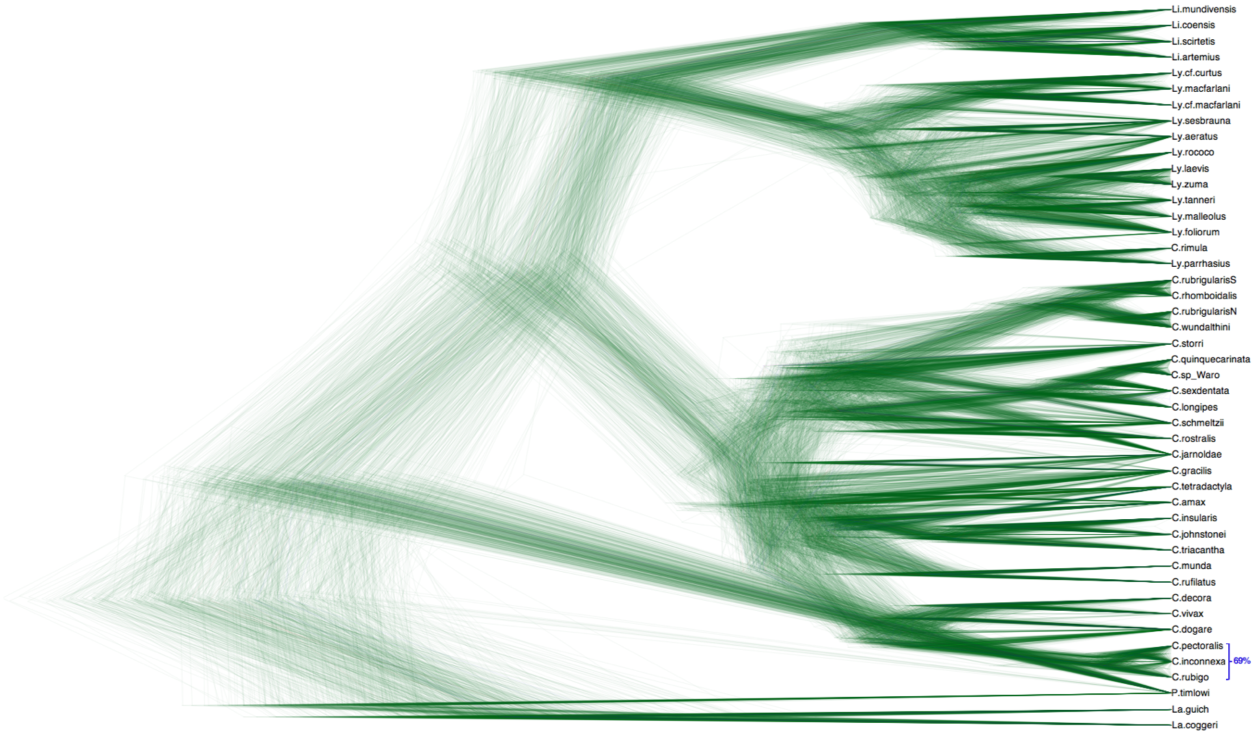


Figure S16: **Cloudogram of species tree samples from UMCMC.** There are 900 samples shown in the above figure. For UMCMC, the 304 loci were separated into 9 subsets and 100 samples per subset were randomly selected from each subset.

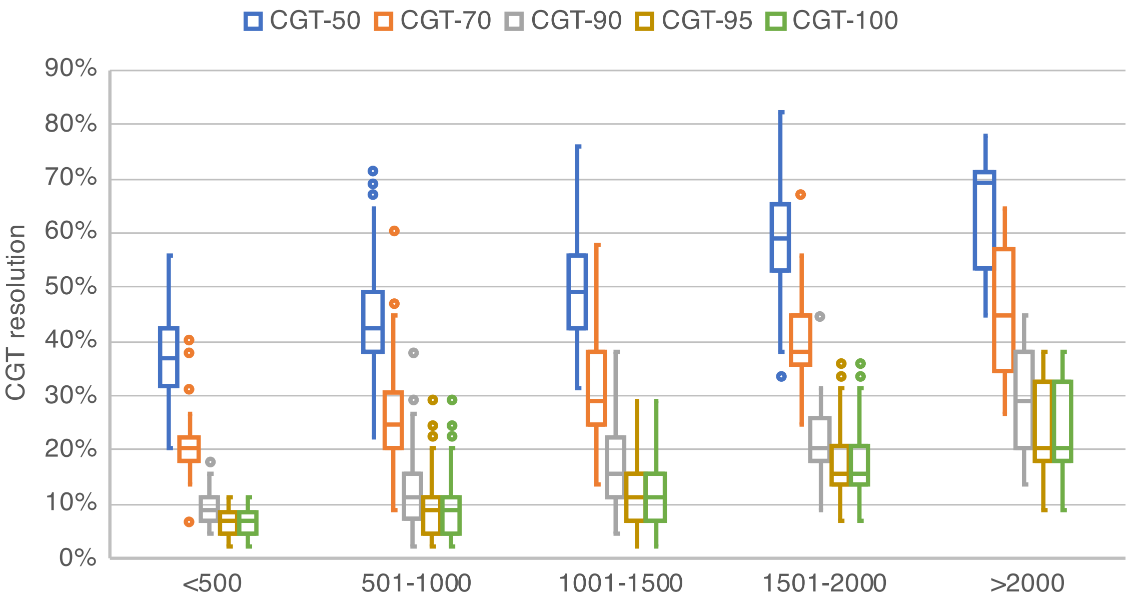


Figure S17: **The proportion of resolved internal nodes for each consensus gene tree in the Australian skink dataset.** Gene trees were partitioned by alignment length along the x-axis. The resolution ratio is defined by the proportion of resolved internal nodes in the CGT in all internal nodes in according gene tree. Different colors represent different consensus thresholds. The length of the 304 sequence alignments varies from 240 to 6,534 sites. In general, as the length of sequence increases the resolution improves.

### S2 Supplementary Tables

Table S1: Species name map between the biological dataset and the paper.

| Species in dataset | Species in the paper |
|---|---|
| Carlia.kimbissp | Carlia.insularis |
| Liburnascincus.cfcoensis | Liburnascincus.artemis |
| Lygisaurus.Melvillesp | Lygisaurus.cf.macfarlani |
| Lygisaurus.novaeguineae | Lygisaurus.cf.curtus |
| Carlia.fusca | Carlia.sp (Waro) |

Table S2: The details of the biological dataset including genus (focal clade), species, tissue, collection and resource library are provided in this table.

| Genus (Focal Clade) | Species | Tissue | Collection | Library |
|---|---|---|---|---|
| Carlia (ECI) | amax | ABTC29892 | MAGNT | SP03_indexing25 |
| Carlia (INS) | insularis | R117953 | Western Australian Museum | AS01_indexing45 |
| Carlia | decora | conx5115 | Queensland Museum | SP07_indexing4 |
| Carlia | dogare | ABTC32199 | Queensland Museum | SP07_indexing5 |
| Carlia | gracilis | CCM0457 | Moritz lab ANU | SP05_indexing14 |
| Carlia | inconnexa | J89138 | Queensland Museum | SP08_indexing7 |
| Carlia (JAR) | jarnoldae | ABTC1107 | Queensland Museum | SP07_indexing8 |
| Carlia | johnstonei | R171237 | Western Australian Museum | AS01_indexing29 |
| Carlia | longipes | ABTC11002 | Australian Museum | SP07_indexing9 |
| Carlia | munda | R131750 | Western Australian Museum | SP04_indexing43 |
| Carlia | pectoralis | ABTC76882 | South Australian Museum | SP07_indexing14 |
| Carlia | quinquecarinata | ABTC102373 | Queensland Museum | SP07_indexing15 |
| Carlia | rhomboidalis | ABTC80487 | South Australian Museum | SP10_indexing20 |
| Carlia | rostralis | A006771 | Queensland Museum | SP09_indexing26 |
| Carlia | rubigo | J89141 | Queensland Museum | SP07_indexing17 |
| Carlia | rubrigularis-N | SS33 | Moritz lab ANU | SP03_indexing6 |
| Carlia | rubrigularis-S | SS46 | Moritz lab ANU | SP02A_indexing7 |
| Carlia (KIM) | rufilatus | CMWA35 | Moritz Lab ANU | SP04_indexing9 |
| Carlia | schmeltzii | ABTC11024 | Australian Museum | SP07_indexing18 |
| Carlia | sexdentata | ABTC10982 | Australian Museum | SP07_indexing19 |
| Carlia | sp. (Waro) | ABTC44734 | Australian Museum | SP07_indexing6 |
| Carlia | storri | A010492 | Queensland Museum | SP09_indexing40 |
| Carlia (TET) | tetradactyla | ABTC11042 | Australian Museum | SP07_indexing20 |
| Carlia | triacantha | R168590 | Western Australian Museum | AS01_indexing16 |
| Carlia | isostriacantha | R168590 | Western Australian Museum | AS01_indexing16 |
| Carlia | vivax | A006791 | Queensland Museum | SP09_indexing3 |
| Carlia | wundalthini | conx5328 | Hoskin collection | SP09_indexing28 |
| Lampropholis | coggeri | SS60 | Moritz lab ANU | SP02A_indexing4 |
| Lampropholis | guichenoti | ABTC12335 | South Australian Museum | SP07_indexing28 |
| Liburnascincus | artemis | conx5371 | Hoskin collection | SP09_indexing29 |
| Liburnascincus | coensis | A004566 | Queensland Museum | SP09_indexing17 |
| Liburnascincus | mundivensis | ABTC10839 | Australian Museum | SP07_indexing11 |
| Liburnascincus | scirtetis | A002000 | Queensland Museum | SP09_indexing19 |
| Lygisaurus | aeratus | ABTC10855 | Australian Museum | SP09_indexing5 |
| Lygisaurus | cf. curtus | ABTC46164 | Australian Museum | SP08_indexing12 |
| Lygisaurus | cf. macfarlani | ABTC30000 | MAGNT | SP07_indexing30 |
| Lygisaurus | macfarlani | conx5614 | Hoskin collection | SP09_indexing39 |
| Lygisaurus | foliorum | ABTC72910 | South Australian Museum | SP08_indexing29 |
| Lygisaurus | laevis | A000355 | Queensland Museum | SP09_indexing11 |
| Lygisaurus | malleolus | A006770 | Queensland Museum | SP09_indexing13 |
| Lygisaurus | parrhasius | ABTC31978 | Queensland Museum | SP08_indexing13 |
| Carlia | rimula | A004595 | Queensland Museum | SP09_indexing2 |
| Lygisaurus | rococo | LR7 | A. Pintor collection | SP07_indexing31 |
| Lygisaurus | sesbrauna | A004711 | Queensland Museum | SP09_indexing15 |
| Lygisaurus | tanneri | A004762 | Queensland Museum | SP09_indexing16 |
| Lygisaurus | zuma | A000129 | Queensland Museum | SP09_indexing8 |
| Pygmaeascincus | timlowi | A001585 | Queensland Museum | SP09_indexing21 |

### S3 Instruction of External Tools

#### S3.1 Generating simulated datasets

For all simulated datasets, we use dendropy (Sukumaran and Holder, 2010) to obtain random species tree given a specific number of taxa. Note that both the topology and the branch lengths of the species tree are randomly generated while the topology is discrete and the branch lengths are positive continuous number. Once the species trees are created, we generate 200 gene trees by MS (Hudson, 2002) for each random species tree. Sequences data are generated by Seq-gen (Rambaut and Grass, 1997) under Jukes-Cantor model. We get the constraint gene tree for each gene locus by bootstrapping from the sequences by RAxML (Stamatakis, 2014). To explore the relationship between the performance of CGT sampler and the consensus threshold, we used 5 consensus thresholds: 50, 70, 90, 95 and 100.

##### S3.1.1 Generating random species trees

To generate species trees with fixed number of taxa or leaves under a birth-death model, we use a Python library dendropy. The following command generates a birth-death tree with 16 taxa. An example of the generated tree is shown in Figure S18
**treesim.birth_death_tree(0.1, 0.025, num_extant_tips = 16, gsa_ntax = 160)**

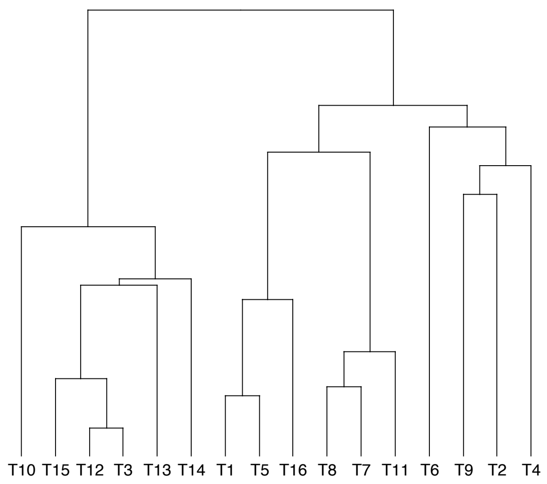


Figure S18: A generated birth-death tree with 16 taxa/leaves.

##### S3.1.2 Generating random gene trees

We use **ms** to generate gene trees given a species tree. We provide all **ms** commands we used are available online: `https://drive.google.com/file/d/1T56Hz3tMCkMU0qXs8-CftFJwsooKwBao/view?usp=sharing`. As an example, the following command generates 200 gene trees given the 16 taxon species tree in Figure S18:

msdir/ms 16 200 -T -I 16 1 1 1 1 1 1 1 1 1 1 1 1 1 1 1 1 -ej 0.311850462472261674 3 -ej 0.6788226115488613 8 7 -ej 0.7788627666563492 11 10 -ej 0.8637613055146363 3 2 -ej 1.1658150944020498 12 10 -ej 1.752813957683974 9 7 -ej 1.9113562469046457 5 2 -ej 1.9900619294961446 6 2 -ej 2.5699665737397805 2 1 -ej 2.931181733823259 15 14 -ej 3.2581218506827128 16 14 -ej 3.40509064102649 10 7 -ej 3.691367778151333 14 13 -ej 3.929597912051401 13 7 -ej 5.00000000000037 7 1

##### S3.1.3 Generating sequence data

After we obtain the phylogenetic tree of each gene locus, we use Seq-gen to generate DNA sequences under Jukes-Cantor model. The length of each sequence alignment is 1,000 under two population mutation rates **0.01** and **0.001** according to "old divergence times" and "young divergence times" in Table 1. The Seq-gen commands are:
**Old divergence times:**
Seq-Gen-1.3.4/source/seq-gen -mHKY -s0.01 -l 1000 -on < input.tree > sequence.nex
**Young divergence times:**
Seq-Gen-1.3.4/source/seq-gen -mHKY -s0.001 -l 1000 -on < input.tree > sequence.nex

#### S3.2 Deriving constraint gene trees

We used RAxML to get the constraint trees. Firstly, we generated 50 bootstrap trees given an alignment. Then, we estimated the constraint trees given a specific consensus threshold. The following two commands show how to derive a constraint tree when the consensus threshold is 50.

raxmlHPC-PTHREADS -m GTRGAMMA -p 12345 -# 50 -s 0/dna.phy -n T0
raxmlHPC-PTHREADS -m GTRGAMMA -J T_50 -z RAxML_bootstrap.T0 -n T1

### References


Hudson, R. R. 2002. Generating samples under a Wright–Fisher neutral model of genetic variation. *Bioinformatics*, 18(2): 337–338.

Kuhner, M. K. and Felsenstein, J. 1994. A simulation comparison of phylogeny algorithms under equal and unequal evolutionary rates. *Molecular Biology and Evolution*, 11(3): 459–468.

Rambaut, A. and Grass, N. C. 1997. Seq-Gen: an application for the Monte Carlo simulation of DNA sequence evolution along phylogenetic trees. *Bioinformatics*, 13(3): 235–238.

St. John, K. 2017. Review Paper: The Shape of Phylogenetic Treespace. *Systematic Biology*, 66(1): e83–e94.

Stamatakis, A. 2014. RAxML version 8: a tool for phylogenetic analysis and post-analysis of large phylogenies. *Bioinformatics*, 30(9): 1312–1313.

Sukumaran, J. and Holder, M. T. 2010. DendroPy: a Python library for phylogenetic computing. *Bioinformatics*, 26(12): 1569–1571.
